# Enhanced Zika Virus Suppression by Coupling Ribavirin with Biodegradable Drug Delivery

**DOI:** 10.1101/2025.11.10.687694

**Authors:** J Rodriguez, A Waterman, MR Blahove, JA Saviskas, BG Santos-Villalobos, MA Wallace, JR Carter

## Abstract

Advancement and implementation of nanoparticle-based systems for enhanced drug delivery have become essential for the suppression of viral pathogens due to lack of viable treatment options. This study focuses on characterization and application of mPEG-PCL co-polymers for development of pH-responsive, biodegradable micellar nanoparticles (MNPs) for the delivery of the antiviral drug ribavirin against Zika virus. Synthesis of mPEG-PCL diblock copolymers was confirmed through FTIR and ^1^H NMR, which validated the formation of ester functional groups and accurate structure of copolymers. mPEG-PCL MNPs, loaded with ribavirin, exhibited a mean diameter of approximately 34.29 ± 5.214 nm and a zeta potential of −3.28 ± 0.718 mV, suitable for evading immune detection and facilitating cellular entry. The ribavirin-loaded mPEG-PCL MNPs demonstrated pH-dependent release, with approximately 88% of ribavirin released at pH 5.5, compared to 20% at pH 7.4. This pH-responsive release is crucial for targeted drug delivery within the endosomal pathway. In vitro studies using JEG-3 cells infected with Zika virus showed that ribavirin-loaded mPEG-PCL MNPs achieved an EC50 of 0.22nM, significantly enhancing drug efficacy compared to unencapsulated ribavirin, which required micromolar concentrations to achieve similar effects. The MTT assay results indicated minimal cytotoxicity of the ribavirin-loaded mPEG-PCL MNPs, with approximately 80% cell viability at the highest concentration evaluated. Confocal microscopy and RT-PCR analysis further confirmed the efficient cellular uptake and potent antiviral activity of the ribavirin-loaded MNPs. These findings highlight the potential of mPEG-PCL MNPs as an effective delivery system for broad-spectrum antivirals like ribavirin, enhancing their therapeutic efficacy while minimizing cytotoxicity.

## Introduction

As of this publication, no FDA-approved vaccines or antiviral treatments are available to address many human viral pathogens that impact everyday life. Consequently, patients are left with supportive care as the sole option for managing these infections, which are scarce in resource-limited regions (1, 2). Scientists continue to investigate ribavirin as a viable therapeutic option for various viral infections (3–5) Biocompatible co-polymers are synthesized from monomers selected based on their properties, enhancing the efficacy of drug delivery systems. Key considerations include pH dependence, improved drug solubility, and effective dissociation characteristics (6, 7). These monomers are systematically organized into structures known as copolymers, enabling the investigation of innovative biomedical applications, including drug delivery. This is due to the amphiphilic self-assembly of copolymers into polymeric micellar formations under specific concentrations and experimental conditions (8). By carefully choosing polymers that exhibit characteristics such as pH-induced dissociation into monomeric components, hydrophobicity, and thermal self-assembly, it is possible to assemble novel monomers and copolymers into customizable systems. These advancements can enhance selectivity while reducing toxicity (9, 10).

The application of biomedically friendly co-polymers has captured scientist’s interest as a consequence of their ubiquitous implementation and characteristics that favor biodegradability, significantly increasing pharmacological applications (11). Biodegradable polymers, such as those described in this article, undergo rapid breakdown, absorption, and elimination from the body following intracellular delivery, making co-polymers highly advantageous for biomedical applications (12). Furthermore, the degradation products of these polymers are non-toxic, easily metabolized, and freely excreted, positioning biodegradable co-block polymers as a prudent option for a wide range of environmentally sustainable and biomedical uses (12).

The diblock mPEG-PCL is a promising co-polymer for biocompatible materials, particularly in the context of anticancer compound delivery. Its advantageous properties include pH-induced dissociation, thermal self-assembly, and hydrophobic characteristics, making mPEG-PCL copolymers ideal for biomedical applications (13, 14). The co-polymer diblock mPEG-PCL is designed from the two constituent segments that combine to create a singular diblock strand: polyethylene glycol and poly(ɛ-caprolactone). The patterns of polymer segments are frequently abbreviated using alphabetic notations, which assign specific letters to each corresponding monomer unit.

Biodegradable co-polymeric nanoparticles, comprising segmented biocompatible molecules, have garnered significant interest among researchers due to their potential to form micelles in aqueous environments. These structures are optimized to enhance drug-release properties under given conditions (15). Di-block, co-polymer-based micelles possess two distinct components: the outer shell and the inner core. Research indicates that drug-loaded micelles featuring a hydrophilic outer compartment, such as those constructed from mPEG-PCL, exhibit several advantages within the body. These benefits include prolonged circulation of encapsulated drugs, enhanced stability *in vitro*, and reduced macrophage recognition following intravenous administration (16). Co-polymer nanoparticles synthesized from mPEG-PCL establish a hydrophilic compartment that facilitates the loading of polar molecules (17). Drug delivery system development can be enhanced by designing a cavity for drug loading, which effectively addresses common challenges in drug development. This approach facilitates the achievement of adequate water solubility for intravenous administration without necessitating modifications to the drug’s chemical structure (18, 19). Linking the established pH stimulation of these nanoparticles may effectively target critical intracellular microdomain pH compartments that are critical for viral replication.

Ribavirin, a guanosine analog, has established itself as a primary antiviral agent over an extended period, owing to its extensive spectrum of activity (20). The mechanism of ribavirin-mediated inhibition is focused on its capacity to act as a mutagen, thereby enhancing the frequency of errors during the replication of viral RNA (vRNA) (21). Ribavirin, upon administration, is integrated into vRNA where it exerts a direct inhibitory effect on viral RNA synthesis. This process induces mutations within the viral genome, ultimately resulting in the inhibition of vRNA translation or causing premature termination of vRNA elongation (22). Ribavirin is recognized as a broad-spectrum antiviral agent with demonstrated efficacy in laboratory settings for the suppression of various viral infections. Its effectiveness extends beyond hepatitis C to include notable viruses such as Lassa fever, RSV, dengue virus, and Zika virus, among others (20). Due to concerns related to cytotoxicity, ribavirin is presently U.S. FDA approved exclusively for the treatment of hepatitis C (23).

As foreshadowed by the preceding, the research presented in this article addresses existing limitations in the antiviral field by developing a novel drug delivery strategy utilizing mPEG-PCL copolymer micellar nanoparticles (MNPs) in conjunction with the antiviral agent ribavirin. Ribavirin was encapsulated within pH-responsive, biodegradable MNPs composed of mPEG-PCL diblock copolymers which demonstrated effective ribavirin release at a pH of 5.5. When tested on cultured choriocarcinoma cells (JEG-3 cells), these nanoparticles exhibited enhanced ribavirin uptake compared to unencapsulated ribavirin, while significantly reducing cytotoxicity, as evidenced by MTT assays. Moreover, activity assays (PRA, RT-PCR, and IFA) demonstrate antiviral effectiveness against Zika viruses leading to full suppression of this mosquito and human-borne virus. The research substantiates our hypothesis regarding the potential efficacy of antiviral-loaded mPEG-PCL MNPs as a superior antiviral strategy against viral pathogens, offering a cost-effective solution that is particularly advantageous for resource-limited regions globally.

## Methods

### Reagents

JEG-3 cells(24) and Vero-6 cells(25) were received from the American Type Culture Collection. These cells were and maintained in Dulbecco’s Modified Eagle Medium (DMEM(26)), Invitrogen-Gibco) augmented antibiotics streptomycin (100 μU/ml) and penicillin G (100U/ml), both from Invitrogen-Gibco, and 10% FBS (Atlanta Biologicals). Cells were propagated in a 37°C/5% CO_2_ incubator environment and sub-cultured every 4 days.

### Synthesis of the diblock polymer, mPEG-PCL

Reagents utilized for AB diblock copolymer synthesis include polyethylene glycol methyl ether and methoxy polyethylene glycol (mPEG, M_n_ 5,000), were acquired from Sigma-Aldrich. Additionally, ε-caprolactone and stannous octoate were procured from Alfa Aesar. Dichloromethane and hexane were obtained from Fisher Chemical, while diethyl ether was acquired from VWR.

### Synthesis of capped A monomer

1,4-dioxane was obtained from Fisher Chemical, 4-(Dimethylamino)pyridine) was obtained from Alfa Aesar, and Succinic anhydride was attained from ACROS Organic.

### mPEG-PCL-OH Synthesis

First, ε-caprolactone was linearized and polymerized by ring-opening polymerization (ROP), with methoxy polyethylene glycol functioning as an initiator with stannous octoate functioning as a catalyst. A mixture containing 4 mL of ε-caprolactone, 100 µL of stannous octoate, and 2 grams of mPEG, and was refluxed at 130 °C for a duration of 4 hours, all while under a nitrogen environment. Subsequently, the resulting product solution was dissolved in dichloromethane and precipitated with chilled diethyl ether and hexane at a volume/volume ratio of 9:1. Resulting mPEG-PCL-OH polymers were then retrieved by filtration and stored in a desiccator.

Lastly, mPEG-PCL co-polymers were suspended in a 1:1 volume/volume of chilled diethyl ether and hexane, followed by dialysis at room temperature with regenerated cellulose dialysis tubing possessing a molecular weight range of 12,000-14,000 against ultrapure deionized water for 12 hours, incorporating intermittent water changes every 4 hrs. The resulting mixture was subsequently lyophilized and stored at −20°C until analyzed by FT-IR spectroscopy and NMR, or until utilized for the assembly of mPEG-PCL MNP nanoparticles.

### Synthesis of mPEG-PCL Micellar Nanoparticles (MNPs)

The AB diblock polymer, composed of mPEG-PCL, was employed to create blank micelles featuring a void core, specifically non-drug-loaded MNPs (27, 28). To synthesize mPEG-PCL MNPs, mPEG-PCL diblock copolymer was dissolved in ultrapure deionized water at a ratio of 20:1 w/w polymer to water. The solution was then positioned in a sonicating water bath at 50°C/15 minutes to facilitate MNP formation (see Figure 1). Upon completion of this process, mPEG-PCL MNPs were passed through a syringe filter, with 0.2 µm pore size, to ensure nanoparticle formation and to eliminate any excess mPEG-PCL co-polymer. Resulting filtered blank micelles (not encapsulating ribavirin) were analyzed by UV-Vis spectroscopy, DLS and zeta potential (ζ).

**Figure 1:**
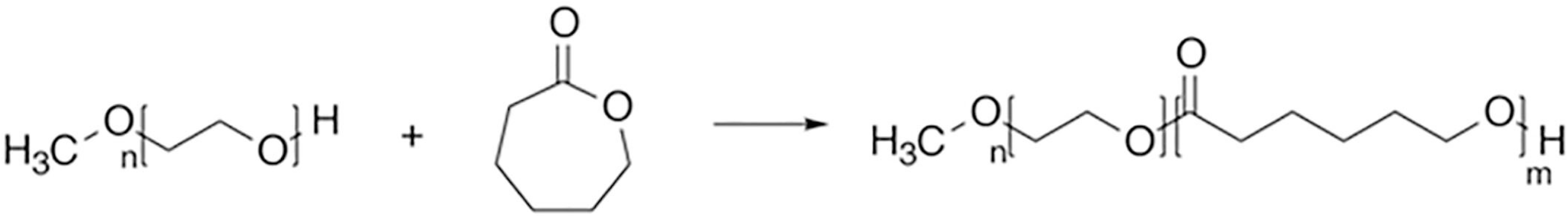
The synthesis of mPEG-PCL co-polymer. The formation of the AB diblock polymer occurs in two stages: First, open ring polymerization of ε-caprolactone (PCL) is initiated using polyethylene glycol (mPEG) as the macroinitiator. Second, stannous octoate is utilized as the catalyst, which is combined with the mPEG-PCL-OH diblock. This process involves modifying the hydroxyl polyethylene glycol into a carboxylic acid, allowing connection to the previously formed diblock. These reactions ultimately yield the completed mPEG-PCL.

### Assembly of Ribavirin-Infused mPEG-PCL MNPs

The required concentration of ribavirin for complete drug loading of our mPEG-PCL micellar nanoparticles (MNPs) was determined based contingently by the topological polar surface area (TPSA) of ribavirin (29), which is a predictive measurement of the polar surface area of a molecule. The TPSA of ribavirin can be computed and evaluated related to drug loading capacity (DLC) of Doxorubicin, which has been extensively investigated in MNP-based drug delivery systems.(30–32) The TPSA of ribavirin was calculated to be 143.73 Å² using Molinspiration Cheminformatics software (Slovensky Grob, Slovak Republic). In comparison, doxorubicin exhibited a TPSA of 206 Å², also determined through the same software. These TPSA values correspond to the DLC 3 mg DOX per 20 mg of mPEG-PCL MNPs, while ribavirin demonstrates a DLC of 4.5 mg ribavirin per 20 mg of mPEG-PCL.

The mixture was subsequently sonicated in an ultrasonic sonicating water bath (Bransonic) at 37°C for 1 hour, maintaining a consistent high pulse setting. The resulting solution was then passed through a syringe filter with 0.2 µm pores to eliminate aggregated ribavirin that failed to load into mPEG-PCL MNPs. The final characterization of the newly ribavirin-loaded mPEG-PCL MNPs was conducted using UV-Vis spectroscopy and DLS to confirm infusion of ribavirin into mPEG-PCL MNPs. The sizes and ζ potential of mPEG MNPs were measured by DLS at 25 °C with a detection angle of 173°.

### Molecular characterization mPEG-PCL copolymers

The structural characteristics of synthesized mPEG-PCL copolymers were evaluated using ^1^H NMR spectroscopy at a frequency of 400 MHz, utilizing a Bruker Avance 500 system from Bruker Biospin GmbH in Rheinstetten, Germany. Molecular weight distribution of mPEG-PCL (Mw/Mn) was determined by gel permeation chromatography (GPC). The system used included a RID-10A refractive index detector and a Shimadzu LC-20AD HPLC pump (Shimadzu, Kyoto, Japan). The polymer mixture was formulated by dissolving mPEG-PCL copolymers in tetrahydrofuran (THF), which also functioned as the mobile phase during GPC analysis. The mobile phase, containing mPEG-PCL copolymers, was then eluted through stationary phase columns of the dimensions 300 mm x 7.8 mm, with a 5 µm particle size (Phenogel™, Phenomenex).

### Scanning electron microscopy (SEM) Analysis of mPEG-PCL MNP Morphology

Characterization of micellar structures and morphologies was conducted using scanning electron microscopy (JEOL JSM-6610LV SEM) to examine the size and shape of the particles. A 0.2% solution of phosphotungstic acid (pH = 7.2) was applied to 300 square mesh copper grids coated with formvar and allowed to air dry for approximately one hour. The particles were then observed under the SEM, which was operated at 30 kV.

### *In vitro* Study of Ribavirin Release from mPEG-PCL MNPs

This protocol wase amended from previously published research (33). Briefly, a 1.5 mL suspension of ribavirin-loaded MNPs was added to 0.5 mL acetate buffer solution (0.1 M, pH 5.5) or 0.5 mL PBS (0.1 M, pH 7.4), and was placed into a 50 kDa dialysis tube. The dialysis tube was then submerged into a 15 mL buffer as is stated for which buffer ribavirin-loaded MNPs have been stated to be submerged in (i.e. acetate or PBS). Following this experimental assembly, filled dialysis tubes were maintained at 37°C, and at the time intervals indicated 1.0 mL of the buffer solution external to the dialysis bag was analyzed by UV-Vis spectroscopy and exchanged with new buffer of equal volume.

### Analysis of Nile Red-Containing mPEG-PCL Micelles for *In Vivo* Tracking of Drug Internalization

Nile red, a fluorescent tracking dye, was incorporated into “blank” mPEG-PCL MNPs to serve as a control for assessing cellular uptake. The loading method for Nile red into mPEG-PCL micelles closely followed the procedure used for ribavirin loading, and as previously reported (34, 35)

### Assessment of Drug Loading Quantitation

Evaluation of drug encapsulation efficiency (DEE) and drug loading capacity (DLC) of our mPEG-PCL MNPs was performed using a standard HPLC with a UV detector at 204 nm. These parameters were calculated based on the following formulas, as previously reported (36):

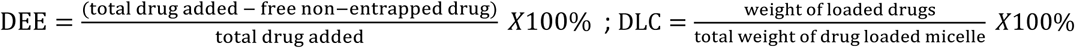

### MTT-Based Cytotoxicity Assay

Ribavirin-encapsulating mPEG-PCL MNPs were assessed for cytotoxicity in JEG-3 choriocarcinoma cells using MTT assays. These assays were performed as previously described (37), but with a change in the concentration of ribavirin used. The ribavirin concentrations in the mPEG-PCL MNPs varied from 1 µM to 12 µM, reflecting the levels typically observed in patients undergoing ribavirin therapy (21, 38). Optical densities were assessed at a wavelength of 492 nm, with a microplate reader (AccuSkanFC, Fisher Scientific). The following formula was used to calculate cell viability:

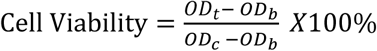

OD_t_ and OD_c_ represent the average optical density (OD) values of surviving cells subjected to an experimental/test group and a control group groups, respectively (39, 40). In contrast, OD_b_ indicates the optical density values of the background in 96-well plates (n = 5). The resulting graphs were produced using GraphPad Prism software.

### Cellular Uptake and Intracellular Cargo Release

The assessment of JEG-3 cells for the internalization of both free ribavirin and mPEG-PCL-encapsulated ribavirin was conducted using UV-Vis spectroscopy to evaluate cellular uptake. A ribavirin stock solution was prepared by dissolving ribavirin (10 mg) in 10 ml DMEM, resulting in a concentration of 1 mg/ml, and was subsequently added to cell culture medium, achieving concentrations ranging from 1 µM to 12 µM per 1 × 10^6^ cells in each well. The cellular uptake of ribavirin and ribavirin-encapsulated mPEG-PCL nanoparticles was analyzed in JEG-3 cells cultured in six-well plates, each containing 1 × 10^6^ cells. Simultaneously, but in separate wells, JEG-3 cells were treated with ribavirin-loaded mPEG-PCL at the same concentration range. Cell culture supernatants were collected two hours post-treatment and analyzed via UV-Vis spectroscopy at 270 nm.

To assess intracellular drug/cargo release from the mPEG-PCL MNPs, Nile Red-loaded mPEG-PCL MNPs (50 ng/ml, based on Nile Red content) and free Nile Red (50 ng/ml) were incubated with JEG-3 cells. The internalization of Nile Red was observed using confocal laser scanning microscopy at an excitation wavelength of 588 nm. Following a 30-minute incubation with the MNPs, cells were washed twice with cold PBS. Subsequently, 100 µL of DAPI (Vectashield, Vector Laboratories, Burlingame, California, USA) was added for nuclear staining, and the cells were examined using LSCM (Zeiss, Germany).

### RT-PCR The Detection of Zika virus RNA

RT-PCR confirmation of ribavirin-loaded mPEG-PCL MNPs was performed as previously described with a few exceptions. JEG-3 cells were cultured at a density of 5 × 10^6^ cells/ml in 6-well plates and subsequently exposed to Zika virus (MOI = 1.0). Following this, the cells were treated with ribavirin-loaded MNPs at concentrations of 0.05 nM and 0.5 nM. RNA isolation, purification, and RT-PCR were performed as previously described (41). The inability to detect Zika virus RNA was interpreted as a sign of virus suppression, corroborating the results obtained from plaque assays. The final validation of the antiviral efficacy of ribavirin-loaded mPEG-PCL MNPs using RT-qPCR was conducted with the iTaq Universal SYBR Green One-Step Kit, as previously outlined (42). Similarly, the failure to detect Zika virus RNA through this method was also regarded as evidence of virus suppression, reinforcing the findings from the initial RT-PCR and plaque assays.

### 2.13 Foci analysis of Zika viruses

Performed as previously described (43), with the following exceptions. Following infection with Zika virus (MOI = 1), the JEG-3 cells were treated with ribavirin-loaded MNPs at concentrations of 0.05 nM and 0.5 nM. Cells were fixed with cold 10% formalin in PBS for 30 minutes. Following aspiration of the fixing solution, cells were washed 2x with PBS supplemented with 3% FBS. Chamber slides were blocked for 1 hour in PBS containing 3% FBS and 0.001% Tween-20. After aspirating the blocking media, the slides were washed twice with PBS supplemented with 3% FBS and incubated for 1 hour with the paraflavivirus 4G2 antibody (1:200, ATCC). The chamber slides were then washed three times and incubated for an hour with a secondary antibody conjugated to Alexa Fluor 488 (Invitrogen). Lastly, slides were treated with mounting media containing DAPI (Vector Laboratories; Burlington, CA) before analysis by LSCM.

## Results

### Characterization mPEG-PCL Co-polymers

Diblock mPEG-PCL copolymer synthesis was executed using a method that has been previously reported (Figure 1) (44). Proper organization of the mPEG-PCL diblock was ensured by FTIR and ^1^H NMR. FTIR was performed on the AB diblock, as well as “A monomers” (mPEG) as well as the complete (Figure 2) (44). The emergence of a peak approximately at 1700 cm^-1^ suggests the existence of ester functional groups within the PCL segment. Additionally, a smaller peak at 1100 cm^-1^ indicates ether (C-O) and ester (C=O) stretching, thereby confirming the formation of mPEG-PCL, while a peak at 1720 cm^-1^ corresponds to C=O in PCL. Lastly, a peak near 3000 cm^-1^, which appears alongside the wide OH peak, is associated with a CH stretch in mPEG.

**Figure 2.**
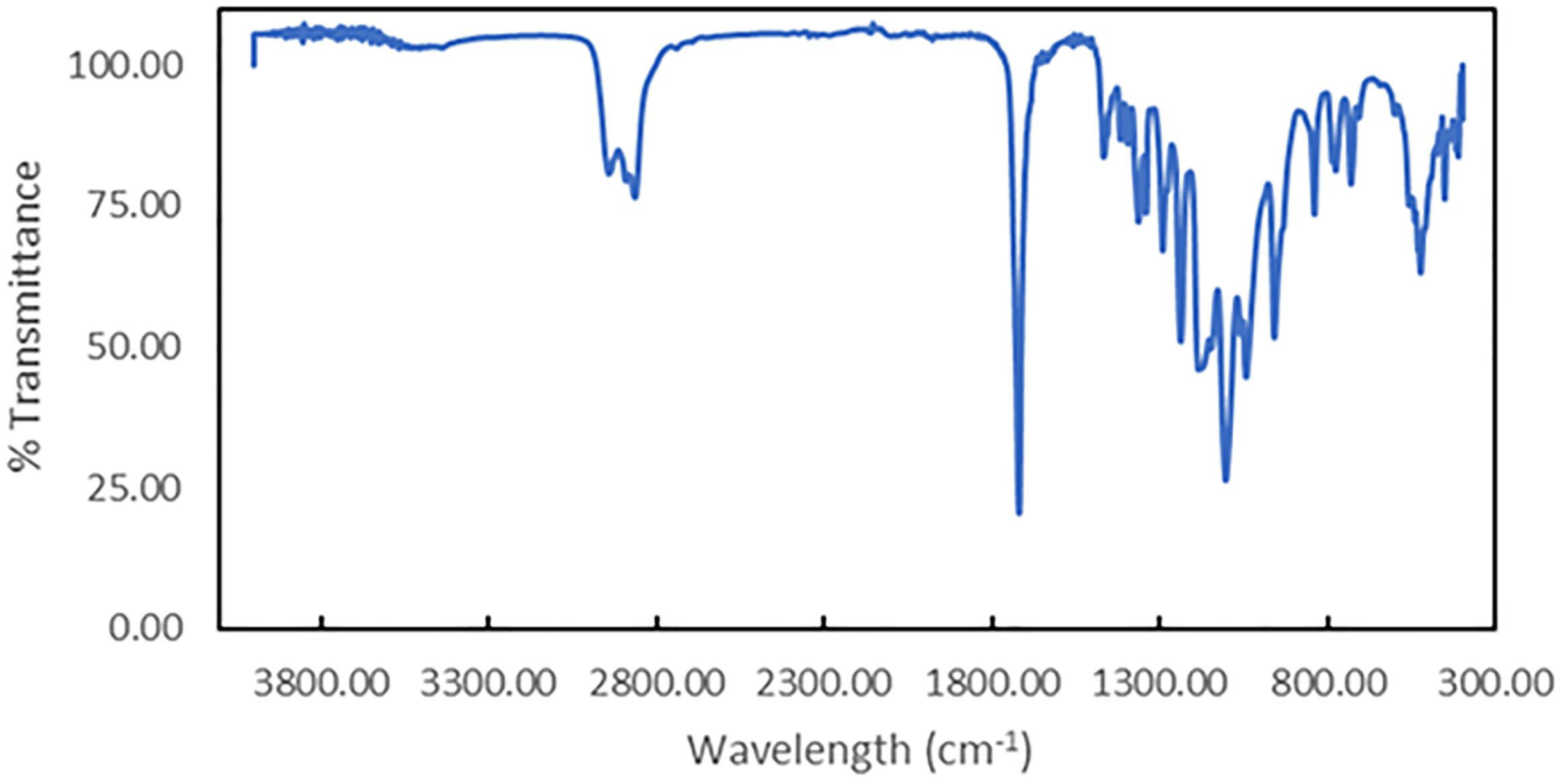
Analysis of the complete diblock polymer, mPEG-PCL by Fourier Transform Infrared (FTIR) spectroscopy.

Proton NMR (^1^H NMR) was employed to further validate the chemical structure of our synthesized mPEG-PCL (Figure 3). The prominent peak observed at 3.59 ppm corresponds to the methylene peak of the methylene glycol (-CH_2_CH_2_O-) unit in mPEG. Additionally, a sharp peak at 2.16 ppm, along with weaker peaks in the ranges of 1.37 ppm and 1.63-1.64 ppm, as well as 2.27-2.32 ppm, were identified as belonging to the methylene protons of the (-OCCH_2_-) groups in the PCL blocks. The signals between 4.02 and 4.04 ppm are attributed to the ethylene protons of the (-O-CH_2_-CH_2_-) units linking mPEG with PCL. Furthermore, the weak peak at 1.80 ppm is associated with the methylene protons of the linker residue dicyclohexylcarbodiimide (DCC). Our NMR findings confirm the accurate structure of the synthesized mPEG-PCL copolymers and align with previously published results.

**Figure 3.**
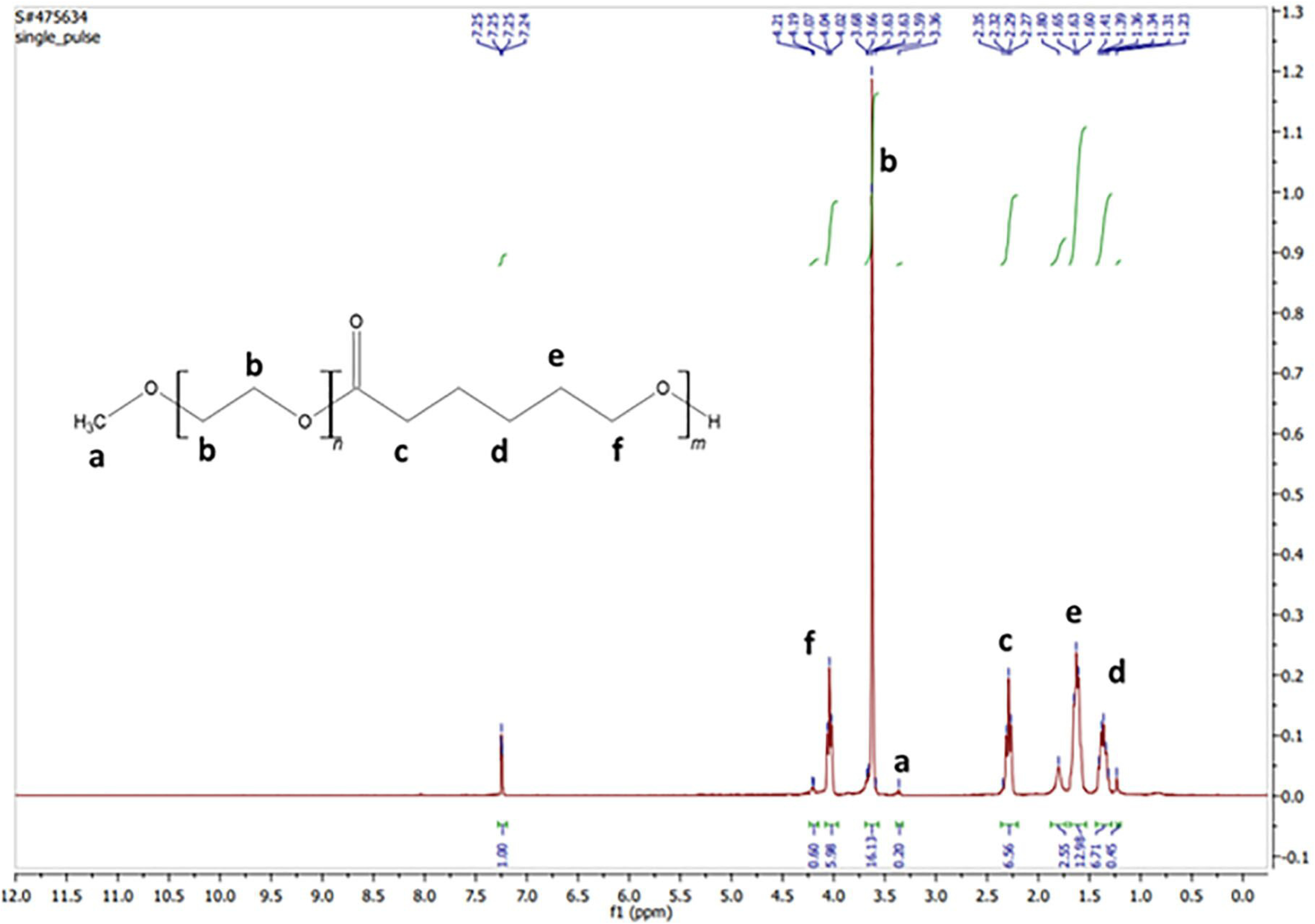
^1^H NMR analysis of mPEG-PCL copolymers in chloroform-D.

The assessment of micellar changes is crucial for understanding the physical properties related to drug delivery. This process involves analyzing the physical characteristics that influence MNP drug loading, which helps predict significant attractive or repulsive forces that may arise when MNPs are further tested *in vitro*. Following mPEG-PCL co-polymer formation into MNPs and subsequent ribavirin loading (Figure 4), surface charge of our mPEG-PCL MNPs was measured using zeta potential instrumentation (Figure 5A), and mean diameter of our ribavirin-loaded mPEG-PCL MNPs in diH_2_O was assessed by dynamic light scattering (DLS; Figure 5B). Mean zeta potential of these MNPs was found to be −3.28 ± 0.718 mV (Figure 5A), while the mean diameter was approximately 34.9 ± 5.214 nm (Figure 5B, Table 1). It is vital the surface charge of our mPEG-PCL MNPs remain within −10 to 10 mV to diminish the possibility of combating significant attractive or repulsive forces during in *vivo* testing, and cardiovascular issues (45). Furthermore, it is critical to maintain the mean diameter of our mPEG-PCL MNPs intended for downstream in vivo studies at less than 100 nm. Smaller particles facilitate quicker cellular entry, and the dynamic light scattering (DLS) instrument will produce a correlation function that correlates with particle size (46). Meaning, nanoparticles of this size can penetrate cells more quickly than larger particles (47). The phenomenon of phagocytosis by macrophages presents an additional compelling rationale. Research indicates that nanoparticles (NPs) exceeding 200 nm in size are phagocytosed by macrophages with greater efficiency compared to those smaller than 200 nm (48). The findings provide valuable insights into an additional potential characteristic of the proposed nanoparticles (NPs) outlined in this proposal: their ability to evade detection by the immune system. Furthermore, the mPEG-PCL MNPs were inspected using scanning electron microscopy (SEM), revealing that mPEG-PCL copolymer-based nanoparticles exhibit a relatively round shape, with a predominantly continuous surface (see Figure 5C). Our SEM analysis also supports the size distribution of the mPEG-PCL-PEG pH-dependent, biodegradable micellar nanoparticles. A summary of the physical properties is presented in Table 1. These results validate the spherical morphology, size, and charge distribution, which are crucial for subsequent therapeutic applications.

**Figure 4:**
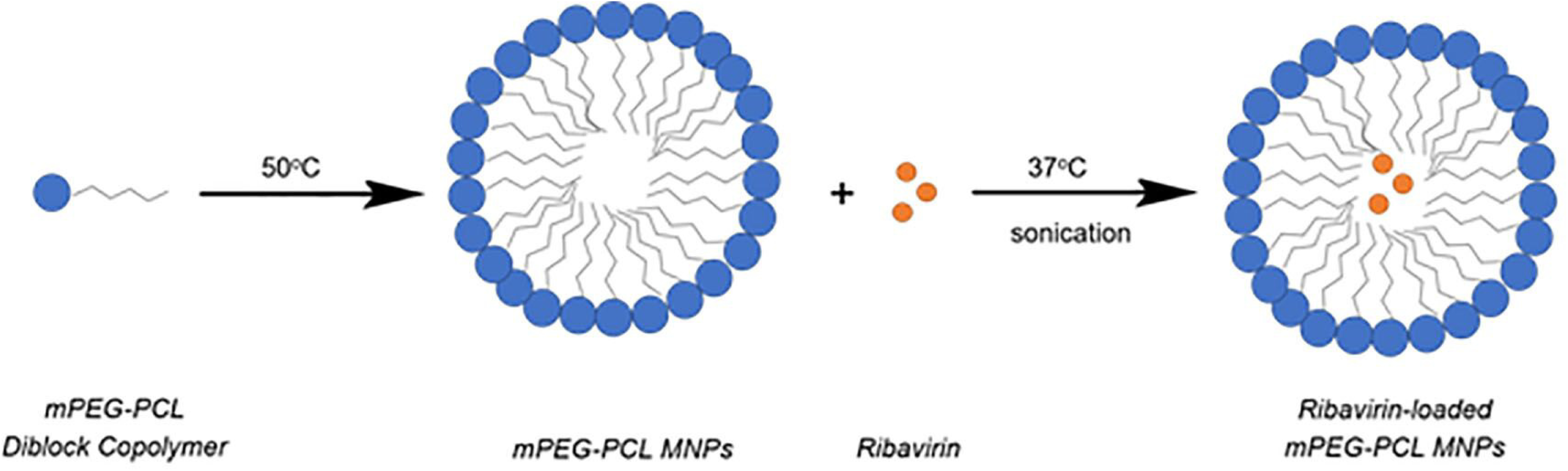
The schematic illustration exhibits the process of creating biodegradable mPEG-PCL diblock ribavirin-loaded mPEG-PCL MNPs. Initially, the mPEG-PCL polymer, is synthesized. Subsequently, micelles are formed at a temperature of 50°C. These mPEG-PCL co-polymer possessing micelles at this stage are referred to as “blanks” in the main text. These newly formed micelles are then loaded with the broad-spectrum antiviral agent, ribavirin, at 37°C through sonication.

**Figure 5.**
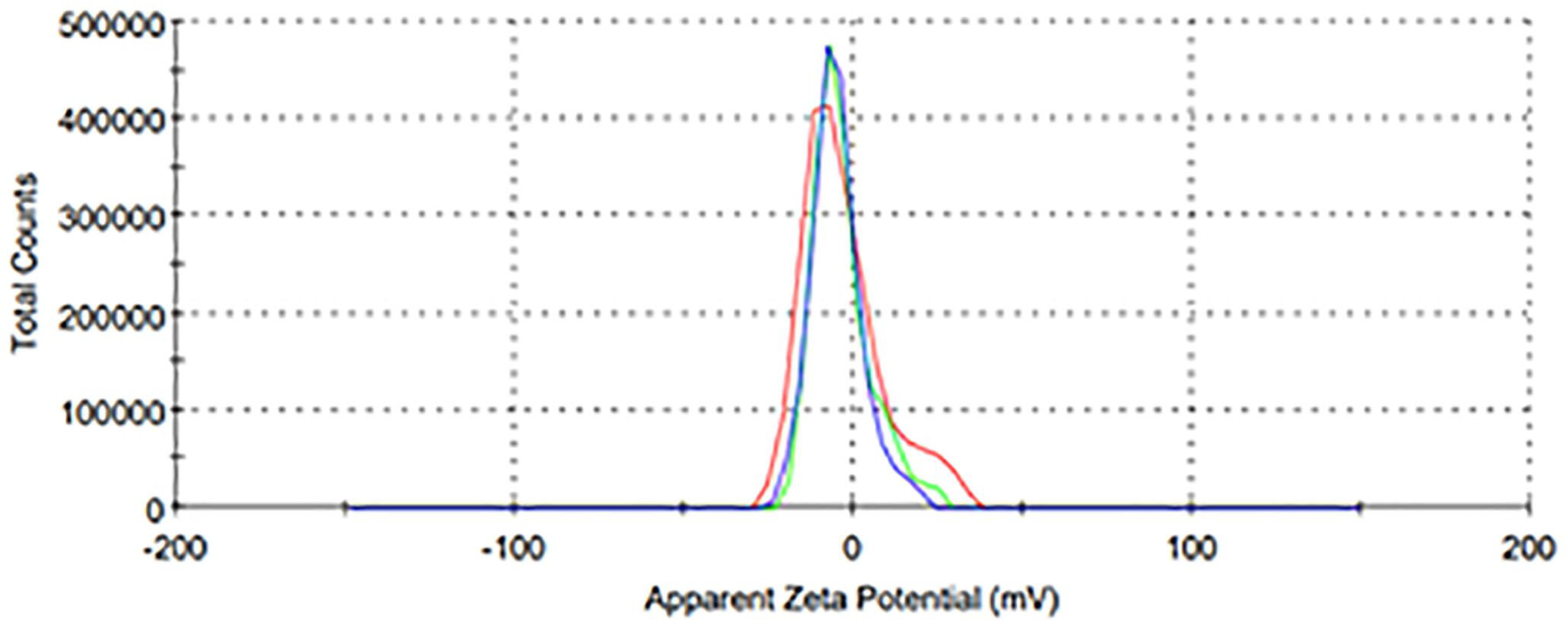
The analysis focuses on the physical characteristics of Ribavirin and Nile Red-loaded mPEG-PCL MNPs. (A) The ζ potential of ribavirin-infused mPEG-PCL MNPs was performed. As outlined in Materials and Methods, the MNPs were suspended in KNO_3_ solution in deionized water to assess their surface charge. (B) The mean diameter of the ribavirin-loaded micellar nanoparticles (MNPs) was determined using DLS, as detailed in Materials and Methods. The DLS spectra illustrates the hydrodynamic size distribution of the synthesized MNPs, with the x-axis representing size in nanometers (nm) and the y-axis indicating intensity in percentage (%). (C) Scanning electron microscopy (SEM) was performed to examine ribavirin-encapsulating nanoparticles made from mPEG-PCL copolymers.

**Table 1:**
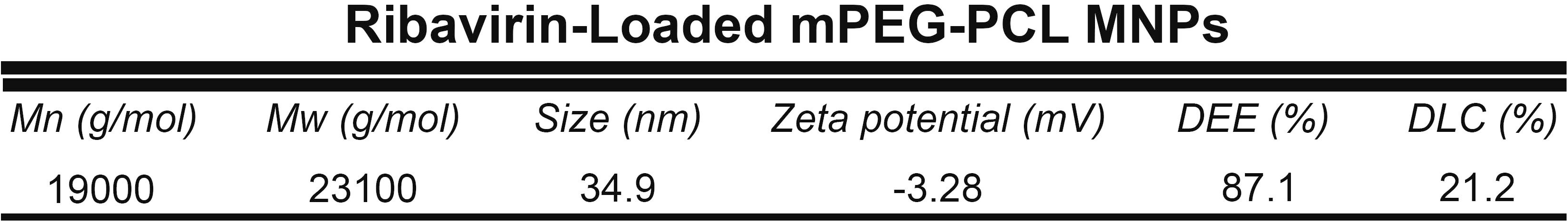
Characteristics of ribavirin-encapsulating mPEG-CL MNPs. Ribavirin-packing, number average molecular weight (M_n_), average molecular weight (M_w_), mean particle size, ζ-potential, and encapsulation efficiency (DEE), of ribavirin-loaded mPEG-PCL MNPs are shown.

### Ribavirin-Release from mPEG-PCL MNPs

mPEG-PCL diblock copolymers comprising MNPs used in this study has been shown to dissociate effectively and entirely at pH 5.5 (49), releasing the encapsulated cargo into the surrounding environment. This finding is compelling, since the pH range of vesicles in the endocytic pathway are between 5.5 (early) and 4.5 (late) (50). As the pH of the endosome decreases during endocytosis, drug cargo (e.g., ribavirin) is released into the endosome and subsequently into the cytosol.

Drug-loaded diblock micelles were synthesized using mPEG-PCL copolymers and the antiviral ribavirin through a process of temperature-induced mPEG-PCL copolymer self-assembly and ribavirin loading (see Figure 4) (27, 36). As assessed by HPLC, the loading capacity (DLC) and encapsulation efficiency were found to be DLC = 21.2 ± 0.03% and DEE = 87.1% ± 3.40%, respectively (Table 1).

Release profiles of ribavirin from our mPEG-PCL MNPs in acetate buffer (0.1 M, pH 5.5) or PBS (0.1 M, pH 7.4) solutions, at 37°C (Figure 6D). An initial explosion of ribavirin release within the first 5 minutes of analysis was observed. This reaction progressed to a constant but slowed release for approximately 25 hours. Over the 25-hour period of analysis with pH 7.4 and 5.5, the amount of ribavirin discharged from our mPEG-PCL MNPs was approximately20% (for pH 7.4) and 88% (for pH 5.5) of the total ribavirin concentration. The rapid release of ribavirin from mPEG-PCL MPNs following pH decrease to 5.5 is potentially associated with repulsion of ribavirin molecules from the hydrophobic core of these MNPs, in a pH-dependent fashion. (51–53)

**Figure 6:**
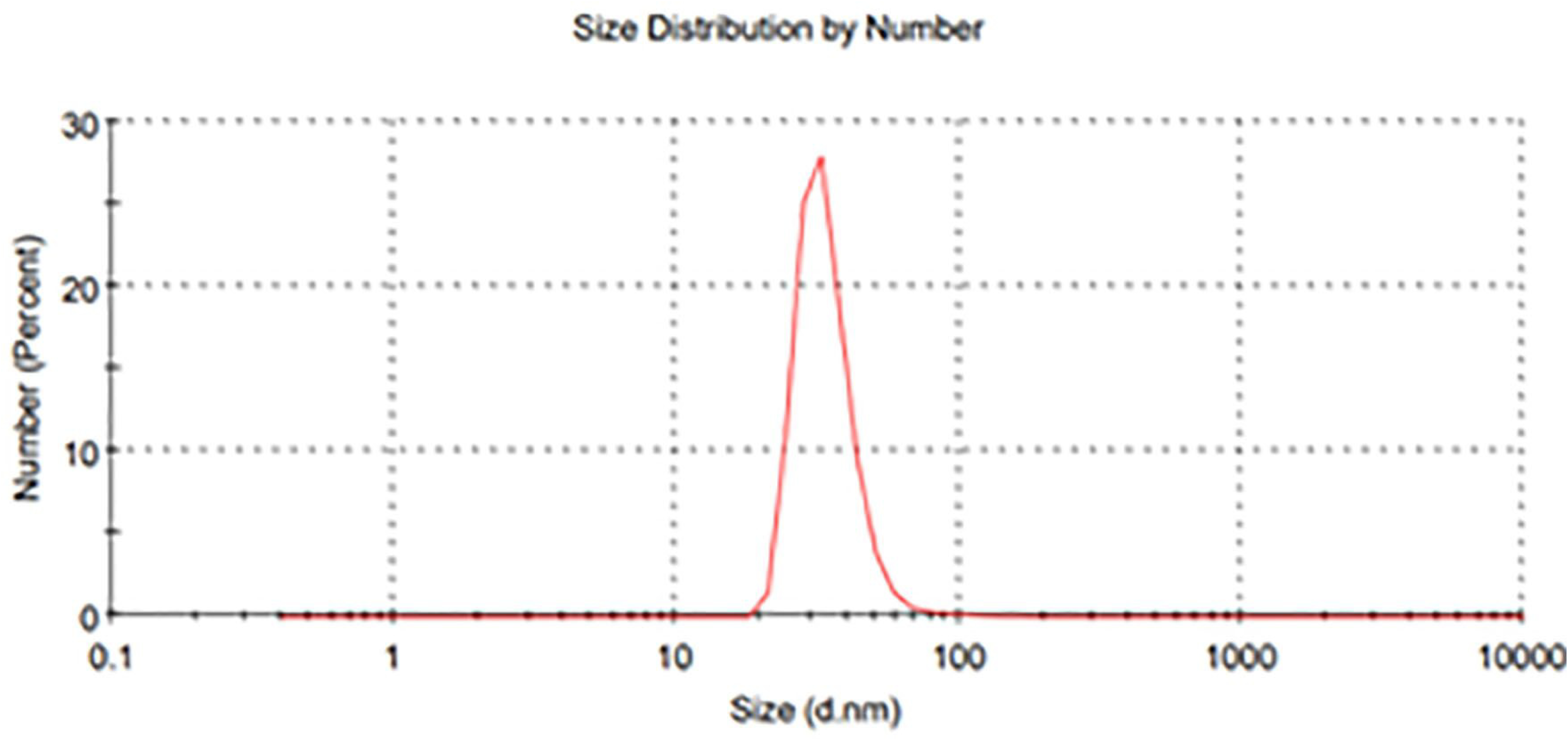
The release profile of ribavirin-infused mPEG-PCL MNPs. Percent of ribavirin is plotted (y-axis) versus the precise time of analysis in acetate (pH 5.5) or phosphate (pH 7.4) buffers. The data shown is the result of 3 independent experiments and was performed as stated in the Experimental Section.

Our results show the most ideal pH-responsive dissociation of mPEG-PCL copolymers occurred at a pH of 5.5, leading to release of approximately 68% more ribavirin from these micelles than at pH 7.4, in a significantly short period of time. Additionally, the release of ribavirin at pH 5.5 is within the range of endosomal pH, target for pH-responsive release (54). Furthermore, the release of ribavirin at this pH level aligns with the typical range of endosomal pH, which is the target for pH-responsive release mechanisms (54).

### Dose Dependence and EC50 Determination by Plaque Reduction Assay (PRA)

The plaque reduction assay was performed in accordance with established protocols outlined in prior research (55). JEG-3 cells in 6-well plates were infected with Zika virus at an MOI of 1.0 virus particles per cell for 1 hour at 37°C. Following Zika virus infection, cells were PBS washed and covered with 1.5 ml of DMEM supplemented with 2% FBS and ribavirin infused MNPs at concentrations ranging from 0.05nM to 0.5nM or free ribavirin at concentrations ranging from 0.1mM to 0.4mM, as indicated in Figure 7. JEG-3 cells (infected and uninfected) were then incubated for 48 hpi at 37°C/5% CO_2_. Next, cell culture fluids were obtained and stored at −80°C for future analysis, which includes quantifying viral titers through plaque reduction assays and detecting Zika virus RNA via RT-PCR, as shown in Figures 7 and 9, respectively.

**Figure 7.**
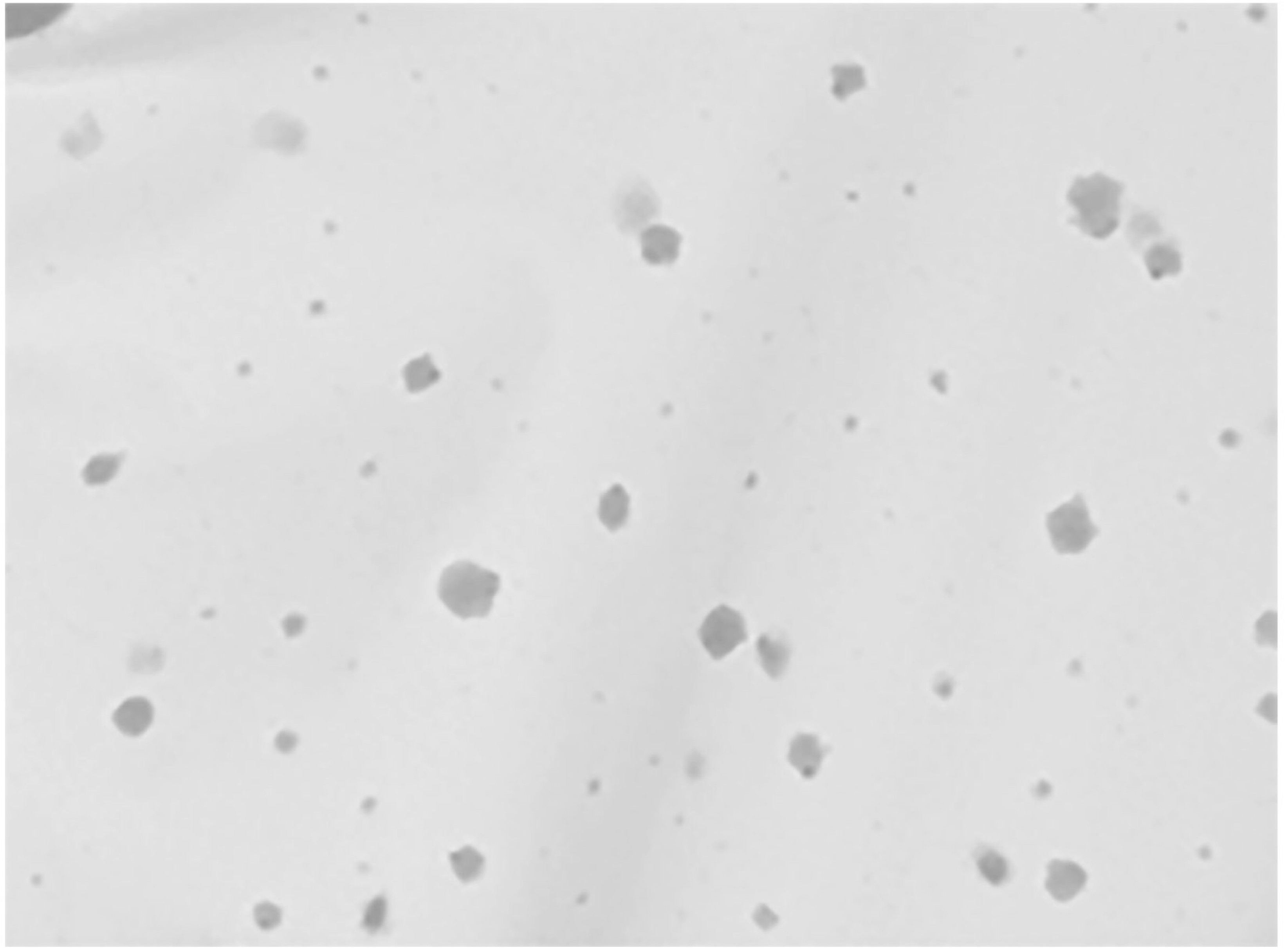
Assessment of ribavirin-loaded mPEG-PCL MNP-mediated suppression by Plaque Reduction Assay. **(A)** Dose-dependent effects of ribavirin-loaded micellar nanoparticles (MNPs) on suppressing Zika viruses. JEG-3 cells were seeded at a density of 1×10^5^ cells per well and subsequently infected with Zika virus at an MOI of 1.0. The cells were treated with ribavirin-encapsulated mPEG-PCL diblock micelles (RM) at the concentrations specified above each well. At 48 hours post-infection (hpi), 10-fold serial dilutions of the culture fluids were prepared and overlaid onto Vero cells, plated at a density of 1×10^4^ in each well of a 24-well plate. After removing the infection media, the cells were overlaid with carboxymethyl cellulose and incubated at 37°C/5% CO_2_ for 5 days. Zika virus-containing plaques were fixed and stained for 20 minutes with 10% formalin and 1% crystal violet solution. The plaques were quantified to assess the efficacy of ribavirin-infused mPEG-PCL micellar nanoparticles in suppressing Zika virus infection. A representative result of the experiment, conducted in triplicate, is presented. **(B)** Quantitation of 3 independent experiments shown in (A). Zika virus titer is indicated on the Y-axis in pfu/ml. Error bars indicate standard deviation of the plaque reduction assay experiments quantitated. Z = Zika virus; D = DMSO; R = unencapsulated ribavirin; RM = ribavirin-infused micellar nanoparticles.

Our plaque reduction assay involved overlaying confluent cultured cells with supernatants infected with Zika virus, utilizing a serial dilution range from 10^1^ to 10^6^ virus particles per milliliter. Cell culture fluids from the initial 48-hour incubation were serially diluted from 10^1^ to 10^6^ virus particles/mL and then overlaid onto confluent Vero cells in 24-well plates. Following the infection of the monolayer with the virus, a 4% carboxymethyl cellulose (CMC) overlay was applied. At five days post-infection, plaques were visible for counting, quantified, and graphed (Figure 7B). As illustrated in Figure 7, ribavirin-loaded mPEG-PCL MNPs protect JEG-3 cells with as little as 0.05nM of ribavirin-loaded mPEG-PCL MNPs added to cells. Furthermore, effective suppression of the Zika virus is achieved with the addition of 0.5nM ribavirin-infused nanoparticles, which is approximately 1000-fold less than the concentration of free ribavirin that elicited a similar result. These results indicate that the application of ribavirin-encapsulated mPEG-PCL MNPs leads to a concentration-dependent reduction in virus titer compared to untreated ZIKV-infected cells (c.f. Z/- to Z/0.05nM and Z/0.5nM). Lastly, it can be hypothesized that a significant level of infection control is due to the presence of a lipid-based micelle, as the addition of the mPEG-PCL “blank” nanoparticle resulted in approximately one log suppression. Collectively, these findings suggest that encapsulating the drug within a pH-dependent, biodegradable micellar nanoparticle like mPEG-PCL results in significantly greater levels of viral suppression compared to administering the drug alone.

Lastly, the results of the plaque reduction assay indicate that encapsulating ribavirin within MNPs made from the biodegradable diblock polymer mPEG-PCL significantly enhances the suppression of the Zika virus in cell culture, achieving an EC50 of 0.22nM. This represents a substantial improvement compared to previously reported EC50 values, which were in the micromolar range for unencapsulated ribavirin (56). The findings indicate that encapsulating antivirals within micellar nanoparticles made from biodegradable copolymers markedly enhances both their solubility and bioavailability.

### MTT Assay results show minimal cytotoxicity in the presence of ribavirin-carrying mPEG-PCL MNPs

MTT assays are utilized to assess cell proliferation, viability, and cytotoxicity, following the protocols outlined in Materials and Methods. Viable JEG-3 cells, which possess active metabolism, are capable of converting MTT into formazan, while dead cells are not able to perform this function and thus result in an absence of signal. Absorbances measured at OD 590 nm correlate with the extent of cell viability when treated with ribavirin-loaded mPEG-PCL MNPs. Assay results indicate that even at the highest concentration tested in our study (12 µM), approximately 80% of JEG-3 cells remained viable (Figure 8, “R-MNP”). In contrast, unencapsulated ribavirin (“Rb”) resulted in approximately 20% cell viability at the same ribavirin concentration. The maximum concentration of 12 µM was selected as it reflects the dosage administered to patients infected with the Hepatitis C virus. These findings demonstrate that the mPEG-PCL encapsulation of ribavirin effectively prevents significant cytotoxicity and cell death.

**Figure 8:**
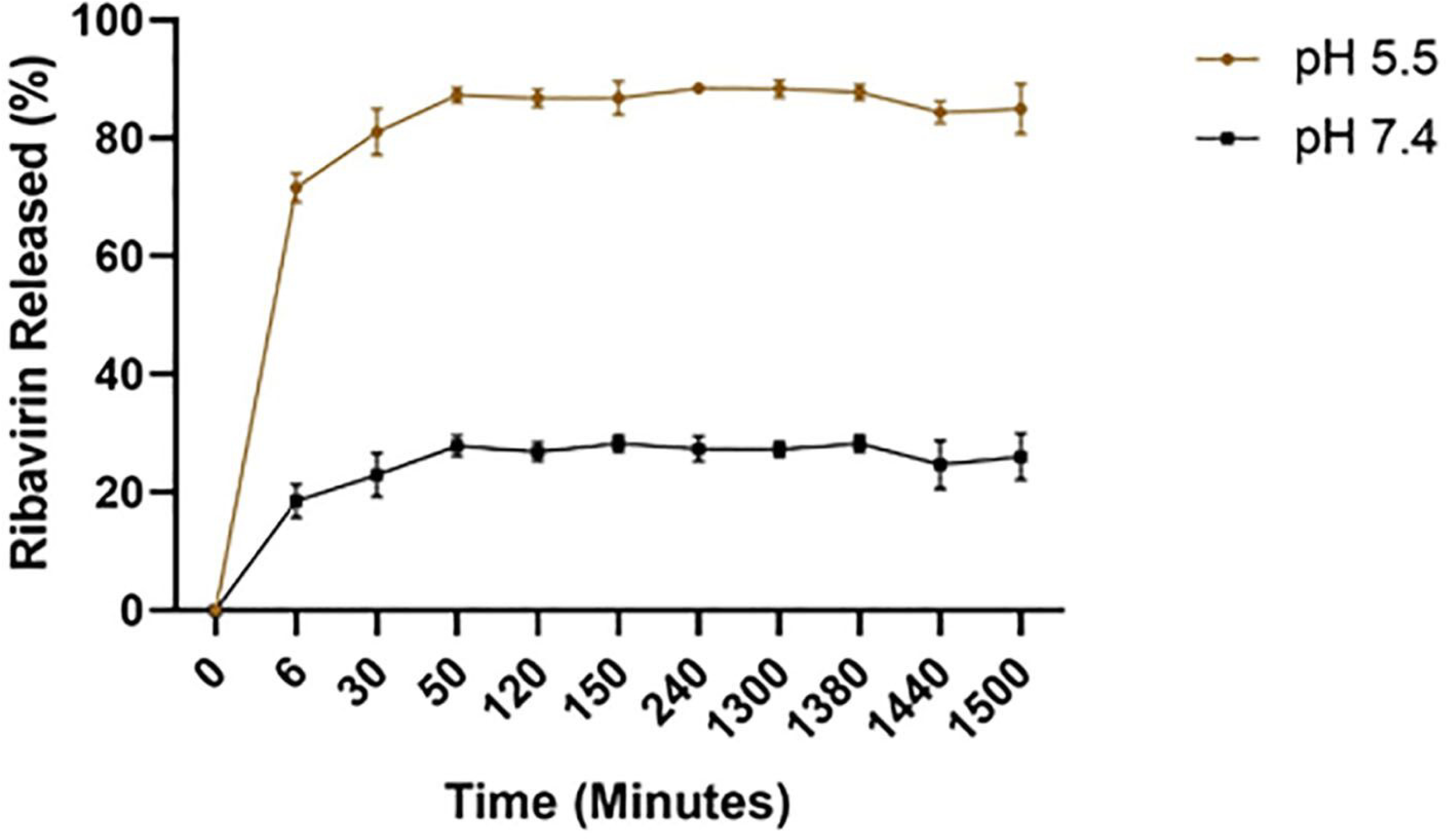
The MTT assay demonstrated that ribavirin-loaded mPEG-PCL micellar nanoparticles did not induce cytotoxicity. The antiviral and cytotoxicity profile of ribavirin was evaluated in JEG-3 cells. Cell viability was measured after adding 12 µM free ribavirin (represented by the black line) or 12 µM ribavirin encapsulated in mPEG-PCL nanoparticles (indicated by the blue line). The concentration of ribavirin encapsulated within the mPEG-PCL nanoparticles was determined using HPLC. The presented results are mean values of three independent experiments, each conducted in triplicate.

### 3.5 Confirmation of Ribavirin-loaded mPEG-PCL MNPs suppression of Zika viruses by RT-PCR

RT-PCR detection of Zika viruses was conducted on Zika virus RNAs extracted from JEG-3 cells and the cell culture media that had been overlaid onto the JEG-3 cells during the infection process. The objective was to assess the effectiveness of ribavirin-loaded nanoparticles in suppressing Zika virion production from both predominantly inactive (cell-derived) and active (viruses that have budded into the cell culture media) sources. Following the assessment of the potential efficacy of ribavirin-loaded mPEG-PCL MNPs through plaque reduction assays (Figure 9), further evaluation of the full suppression achieved by these nanoparticles was performed using RT-PCR.

**Figure 9:**
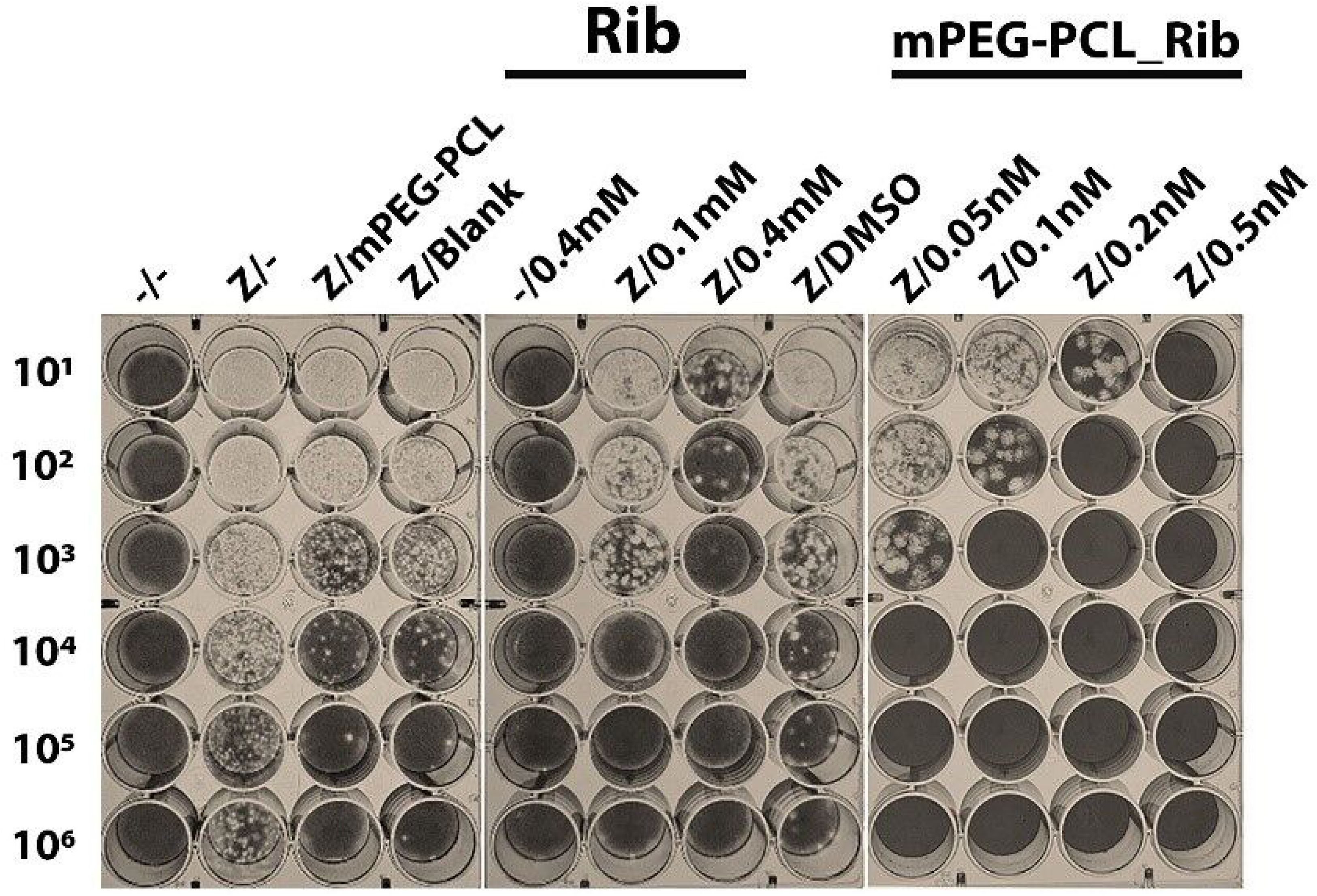
RT-PCR Detection of Zika vRNA Post-Addition of Ribavirin-Loaded mPEG-PCL MNPs. The detection of Zika viruses through RT-PCR was performed, utilizing human placenta choriocarcinoma cells (JEG-3). The cells were cultured at a density of 1×10^5^ in 6-well plates and subsequently infected with the Zika virus at an MOI of 1.0. Next, cells were treated with ribavirin-encapsulated mPEG-PCL polymeric micelles at designated concentrations. The analysis of Zika virus RNAs was performed using either cell culture or cell lysates fluids (Lanes 1 through 6), employing RT-qPCR (A) and RT-PCR (B) methodologies. which successfully amplified a 500 bp fragment, indicating the presence of the virus. Abbreviations: ZIKV = Zika virus; RM = ribavirin-loaded MNPs; ng = nanograms; mL = milliliters.

Placenta carcinoma cells (JEG-3) were infected with Zika virus (M.O.I. = 1.0), while simultaneously treated with ribavirin-loaded mPEG-PCL MNPs at the specified concentrations (Figure 8). Zika virus RNA was extracted from both JEG-3 cells and cell culture fluids, and RT-PCR assays were conducted as per the manufacturer’s instructions (Invitrogen). In comparison to the negative control lanes (lanes 1 and 2), the addition of ribavirin-loaded mPEG-PCL MNPs at a concentration as low as 0.05nM resulted in undetectable levels of Zika virus RNA (lane 6), indicating that full suppression of Zika virus replication was achieved. This outcome was consistent whether analyzing cell culture fluids or JEG-3 cells.

These findings corroborate the results obtained from our plaque reduction assays, demonstrating that ribavirin-loaded mPEG-PCL MNPs can induce complete suppression of Zika virus, utilizing only a fraction of ribavirin compared to previously published results that describe Zika virus suppression by free, unencapsulated ribavirin.

### Analysis of intercellular uptake and cargo release of Nile Red-loaded MNPs through Nile Red uptake assessment

Confirming the effective cellular entry of small molecule-loaded micellar nanoparticles (MNPs), such as ribavirin-loaded mPEG-PCL, is crucial for demonstrating the cellular uptake of our mPEG-PCL drug delivery system. Nile red, a photo-stable and polarity-sensitive fluorescent probe, emits a strong fluorescence at 552 nm and is frequently employed to evaluate cellular uptake (34, 35).

After synthesis of mPEG-PCL copolymers and subsequent formation into MNPs, Nile Red colorimetric molecules were incorporated into the mPEG-PCL nano-micelles. Nile Red was chosen for the *in vitro* analysis of MNP cargo uptake following pH-dependent dissociation of mPEG-PCL MNPs, due to Nile Red’s status as a gold standard in cellular system tracking. The synthesis of Nile Red-loaded mPEG-PCL MNPs was conducted following the same protocol used for ribavirin-loaded mPEG-PCL MNPs, 0.5 mg Nile Red loaded per 20 mg of mPEG-PCL MNPs. Once formed, Nile Red-infused mPEG-PCL MNPs were analyzed with UV-vis spectroscopy in conjunction with DLS to confirm proper loading of the particles. The mean diameter of the Nile Red-imbued mPEG-PCL MNPs was measured at 81.2 ± 4.1nm using DLS, which is considerably greater than that of the ribavirin-loaded mPEG-PCL MNPs, but still within the acceptable range to facilitate cellular uptake. (48) Nile red-loaded mPEG-PCL MNPs not only fulfilled the theoretical criteria but also demonstrated the ability to release drugs in a pH-dependent manner, similar to their ribavirin counterparts.

The intercellular release of Nile Red from Nile Red-loaded MNPs indicates the efficiency of MNPs in delivering drug cargo internally (Figure 10A). To evaluate this, Nile Red-loaded mPEG-PCL MNPs were administered to JEG-3 cells. One hour following Nile Red-loaded-mPEG-PCL MNP addition, cells were washed five times with PBS (pH 7.4) containing 3% FBS and subsequently fixed with 10% formalin. DAPI served as a nuclear stain. The influx of encapsulated Nile Red was compared to that of Nile Red dissolved in DMEM using laser scanning confocal microscopy (LSCM). The results demonstrated that nanoparticle encapsulation facilitated drastically greater levels of cellular uptake compared to the unencapsulated counterparts. Imaging of cellular uptake was conducted one-hour post-incubation. The consumption of Nile Red-loaded MNPs was prompt, with LSCM images revealing that all cells appeared positive for the cytotracker, Nile Red, after just one hour of incubation with mPEG-PCL MNPs. In contrast, cells incubated with free Nile Red dissolved in DMEM did not exhibit any signs of cellular uptake during the same one-hour timeframe. These findings indicate that the cargo contained within mPEG-PCL copolymer MNPs is delivered far more efficiently than without MNP encapsulation.

**Figure 10:**
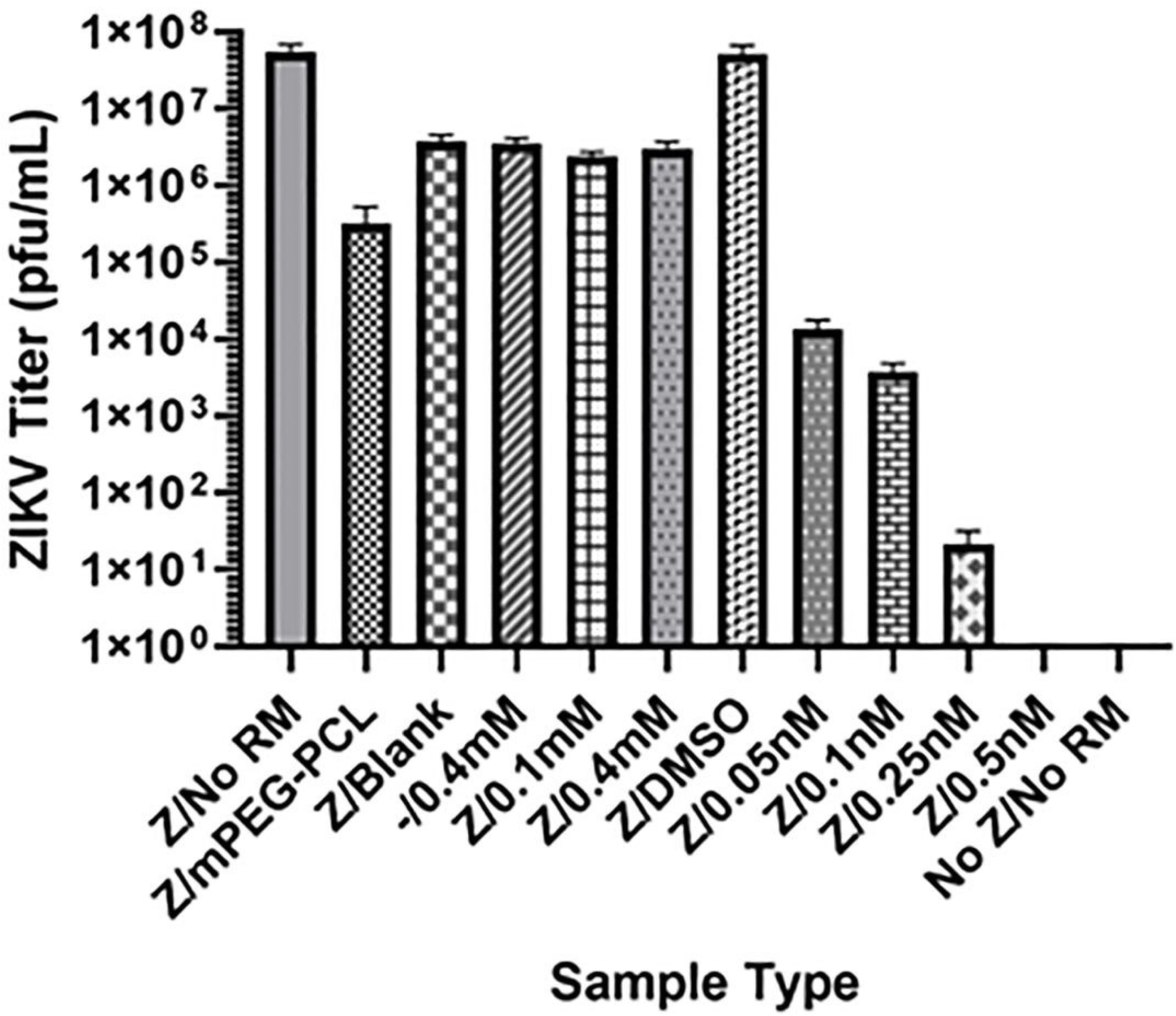
Cellular uptake of Nile Red Ribavirin and encapsulated mPEG-PCL MNPs. A) Confocal analysis of JEG-3 cells, either alone or incubated with Nile Red (50 ng/ml) or mPEG-PCL nanoparticles infused with Nile Red (Nile Red mPEG-PCL), along with DAPI staining for nuclei (blue), revealed the accumulation of nanoparticle-loaded micelles in the cytoplasm after 1 hour of incubation. The merged images demonstrated significant cellular uptake of the polymeric nanoparticles, followed by cargo release. The scale bar represents 100 μm. Nile Red was visualized by laser at 548 nm, and images were captured at 10x magnification. NR-MNP refers to Nile Red-infused mPEG-PCL MNPs. B) The analysis of ribavirin (Rb) and ribavirin-encapsulated mPEG-PCL (R-MNP) involved incubating JEG-3 cells with DMEM containing 12 μM Rb or R-MNP. The ribavirin content was assessed using UV-Vis spectroscopy at 270 nm. A decrease in absorbance indicated ribavirin uptake by JEG-3 cells. The results were compared to controls consisting of cells only and ribavirin only (No Cells).

Human JEG-3 choriocarcinoma cells were cultured into 6-well plates and treated with ribavirin, either encapsulated in mPEG-PCL micelles or in its unencapsulated form, at a final concentration of 12 µM (Figure 10B). This concentration of ribavirin is consistent with levels found in patients undergoing antiviral therapy. Cell culture supernatants were collected two hours after treatment. Notably, cells treated with mPEG-PCL encapsulated ribavirin exhibited a significant increase in drug uptake, approximately fivefold, compared to those receiving unencapsulated ribavirin. These findings support the potential of mPEG-PCL as an effective system for intracellular delivery of broad-spectrum antivirals like ribavirin.

### Immunofluorescence analysis of Zika virus foci

Choriocarcionoma (JEG-3) cells were infected with Zika viruses at an MOI of 1. 0. At 3 dpi, representative LSCM-obtained immunofluorescence images were analyzed (Figure 11). JEG-3 cells infected with Zika displayed a similar pattern of immunofluorescence for the Zika Virus E glycoprotein, which has been shown previously to be localized in the perinuclear and plasma membrane portions of infected cells (57, 58). Immunofluorescence results show a decrease in Zika foci formed is detected in a concentration-dependent manner (0.05nM to 0.5nM), resulting in complete disappearance of Zika foci (i.e., Zika E protein production) and, correlating with the dose dependence assay and RT-PCR results that all suggest ribavirin-loaded mPEG-MNPs possess efficacy in suppressing Zika virus at concentrations fare less than without MNP encapsulation

**Figure 11:**
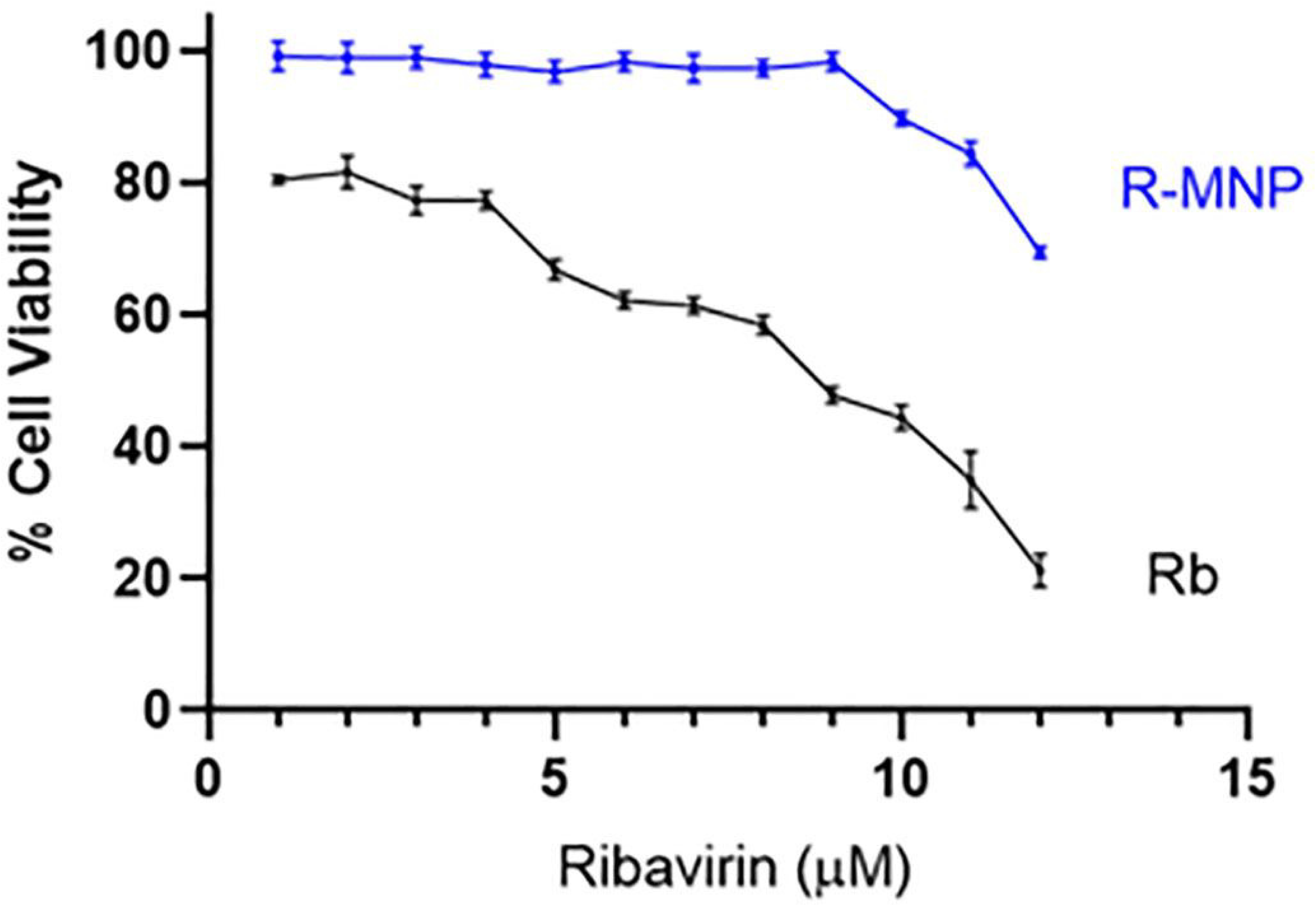
Zika virus foci analysis by Immunofluorescence. At 4 dpi, JEG-3 cells were prepared for immunofluorescence as outlined in the Materials and Methods section. The cells were then incubated with the pan-flavivirus antibody (4G2), followed by incubation with an Alexa-488-conjugated secondary antibody. Fluorescence images were captured at 10x magnification using consistent acquisition parameters. Nuclei were counterstained with DAPI, which appears blue. Rib-mPEG-PCL refers to Ribavirin-infused mPEG-PCL micellar nanoparticles.

## Discussion

Quite recently, nanotechnology has shown promise in fighting infectious diseases. Special tiny particles are being used to develop vaccines, deliver antiviral drugs, and detect pathogens. Nanostructured vaccines, like those for SARS-CoV-2, HPV, hepatitis B, and influenza, have been widely used (59). This gives precedent to the development of nanocarriers as devices to improve intercellular delivery of antiviral compounds.

Ribavirin is a broad-spectrum antiviral medication that has garnered attention for its efficacy against various viral infections (60). Its broad-spectrum activity and established clinical use make it a viable option for treating certain viral infections, particularly in situations where specific antivirals are unavailable or not yet approved. Ribavirin’s antiviral properties have been extensively studied, demonstrating effectiveness against a comprehensive of RNA and DNA viruses in both in vitro and in vivo settings (61). This broad-spectrum nature renders it a potential therapeutic option when the specific viral etiology is unknown or in cases of co-infection with multiple viruses. As we alluded to, until an FDA approved therapy for Zika virus infections emerge, ribavirin should be considered, whether alone or in combination with additional therapies.

Additionally, researchers have investigated ribavirin’s antiviral effects against additional flaviviruses such as dengue virus, albeit with limited efficacy as a singular agent (62). While ribavirin can inhibit dengue virus replication in cell cultures and potentially enhance the efficacy of other antivirals in combination therapy, its prophylactic use has yielded limited success in animal models (63).

In the context of Zika virus, ribavirin has previously demonstrated promise in inhibiting viral replication in vitro and suppressing viremia in animal models, particularly in STAT1-deficient mice. However, further research is necessary to determine its efficacy and safety in humans (56). While no single approved treatment exists for Zika virus (ZIKV) disease, research actively investigates various therapeutic approaches, including the nanoparticle-based drug delivery systems, described in this publication and others. A combination of favipiravir and interferon alpha substantially inhibits ZIKV replication in vitro. Furthermore, nanoparticle-encapsulated ivermectin increases its accumulation in the blood and targets ZIKV’s nonstructural protein 1 (NS1)(64). However, another study showed ivermectin lacks efficacy in preventing Zika-virus-induced lethality (65). Lastly, it has also been shown that even high doses of ivermectin prophylaxis (1.2 mg/kg) for 3 consecutive days) did not prevent Zika virus infection or alter antibody production in rhesus macaques (66). In short, aside from ivermectin lacking FDA approval for use against viral pathogens, due to inconsistent results obtained regarding ivermectin’s efficacy against viral pathogens (e.g. Zika) in cultured cells and animal models, additional research and close monitoring of drug safety is essential.

mPEG-PCL micellar nanoparticles (MNPs) exhibit a favorable profile characterized by biodegradability, biocompatibility, and amphiphilicity, ensuring stability in blood circulation. Furthermore, these MNPs demonstrate a lack of toxicity, immunogenicity, and inflammatory potential, thereby rendering them an exemplary candidate for applications in drug-delivery systems. These systems have shown pH-responsive release of drugs like paclitaxel, making them potentially more effective in targeted delivery. PEG-PCL nanoparticles have been used for targeted delivery and can control drug release by varying the pH (67). Similarly, pyrimethamine-Loaded mPEG-PCL nanoparticles exhibit reduction-responsive release against Toxoplasma gondii under specific conditions (e.g., 10-mM dithiothreitol), which may be beneficial for certain applications. (68).

The copolymers comprising mPEG-PCL MNPs have emerged as favorable materials for drug delivery systems. However, they possess inherent limitations. Our choice of mPEG-PCL is not without its drawbacks. The initial burst release observed (∼6 minutes) is not always ideal drug delivery (Figure 6). However, chemical conjugation of drugs to mPEG-PCL-based MNPs can lead to reduced initial burst release and increased efficiency compared to encapsulation methods (69). This is because conjugation creates a stable, covalent link between the drug and the polymer, which leads to more controlled and sustained drug release over time, reducing premature release and enhancing the drug’s accumulation at the target site. Unfortunately, disadvantages of drug conjugation to mPEG-PCL copolymers and MNPs are numerous, and include premature drug release leading to systemic toxicity, low drug loading due to limited functional groups on the base copolymer, challenges with intracellular drug release and escape from the target cell, and the potential for PEG interference in cellular uptake, all of which require careful balancing of conjugate stability and drug release kinetics for effective and safe drug delivery.

As is the case with all research endeavors, limitations can be found within the data generated. While the data resulting from our proof-of-concept approach involving the effectiveness of ribavirin-encapsulated mPEG-PCL copolymer MNPs in suppressing Zika viruses provides valuable insights, it’s important to acknowledge that limitations exist perpetuating further studies. For example, the small sample size may limit the generalizability of these results (70). Our analysis was limited to JEG-3 cells, a specific cell line, whereas the Zika virus is known to exhibit a broader tropism, infecting a more diverse range of cell types, including astrocytes, neural stem cells, oligodendrocytes, astrocytes, and macrophages (71). Utilizing diverse cell types in culture models for assessing ribavirin-encapsulated mPEG-PCL co-polymer MNPs could yield disparate results. Furthermore, it would be beneficial to investigate the efficacy of ribavirin-encapsulated mPEG-PCL MNPs against a wider array of flaviviruses, such as HCV, dengue, yellow fever, and mosquito-borne alphaviruses like chikungunya. The limited scope of this study could also be extended to encompass a broader range of antiviral compounds. Ribavirin was selected for this investigation due to its broad-spectrum properties and its demonstrated efficacy as a combinatorial therapy with direct-acting antivirals (DAAs), resulting in reduced relapse rates (72). Nonetheless, it would be prudent to evaluate additional antivirals in the context of mPEG-PCL encapsulation to enhance the versatility of this system.

Since we are investigating a previously underexplored facet of ribavirin utilization, specifically within the realm of drug delivery, the shortage of existing literature may impose significant constraints (73). Nevertheless, it is noteworthy that the absence of comprehensive research in this area also presents opportunities for pioneering studies, thereby contributing substantially to the advancement of knowledge within this burgeoning field. In such instances, acknowledging this limitation serves as a pivotal step in identifying knowledge gaps and underscoring the necessity for future investigation into pH-dependent drug delivery and its implications for antiviral therapeutics.

A salient benefit of pH-dependent, biodegradable nanoparticles, exemplified by those formulated from mPEG-PCL copolymers, lies in their distinctive core-shell structure, which confers advantages over other nanoparticulate systems (74). The inner core facilitates the solubilization of hydrophobic pharmaceuticals in aqueous environments, while the mPEG shell provides a protective barrier that mitigates elimination by the mononuclear phagocyte system, thereby extending the duration of blood circulation (75, 76). Furthermore, biodegradable micelles exhibit minimal cytotoxicity, and facilitate renal excretion provided the molecular weight of the constituent polymer chains remains below the threshold value requisite for renal filtration (77, 78).

A mean encapsulation efficiency (EE) of 87.1% ± 3.40% indicating that a substantial portion of the drug has been successfully encapsulated within the nanoparticles. However, this is more a feature of the chemical composition and characteristics of the drug being encapsulated and the co-polymer composition of the pH-dependent micellar nanoparticle, as published data shows an EE range of 98% to 43% (79–81). Our findings further indicate that the encapsulation of antiviral compounds within mPEG-PCL copolymer-based micellar nanoparticles presents a viable drug delivery and antiviral system, distinguished by its efficacy and cost-effectiveness in synthesis and assembly, thereby offering substantial benefits in resource-limited regions where viral outbreaks pose significant challenges, potentially through the provision of a cost-effective and readily synthesizable antiviral platform

Ribavirin encapsulation efficiency and drug loading capacity (Table 1) may also possess limitations that impact overall efficacy of using not only a particular type of pH-dependent nanoparticle (e.g. mPEG-PCL as a drug delivery system, but also the antiviral molecule in question (i.e. ribavirin).

While pH-dependent nanoparticles offer exciting potential for targeted drug delivery, achieving high encapsulation efficiency (EE) and loading capacity (LC) for drugs within these systems can be challenging. For pH-driven encapsulation techniques to be effective, both the core and wall materials must possess pH-dependent solubility, which can be a limiting factor in material selection (82). Some pH-sensitive chemical bonds within the nanoparticle structure, like those based on acetal, hydrazone, orthoester, and ketal functionalities, can cleave rapidly in acidic environments, leading to premature drug release and reduced EE (83). Some cross-linked pH-responsive systems may not fully release their payload within the target pH range, leading to suboptimal delivery and efficacy (84). Encapsulating hydrophobic and large molecular weight drugs within pH-responsive systems can be difficult, resulting in lower EE. Unfortunately, hydrophilic compounds, such as ribavirin, have an issue in EE and LC as well. Highly water-soluble and hydrophilic drugs can be difficult to encapsulate effectively in certain polymeric nanoparticles, leading to drug diffusion out of the nanoparticle during preparation. In some pH-driven nanoparticle formulations, the low volume fraction of the internal aqueous phase compared to the continuous external phase can lead to low encapsulation efficiency. While increasing polymer concentration can improve EE, it can also lead to increased viscosity and potential for aggregation, impacting nanoparticle characteristics (85). While some pH-driven methods aim to avoid organic solvents, their use in other preparation methods can lead to encapsulation of hydrophilic molecules and reduced EE(86).

Premature drug release in non-targeted areas can also be an issue. Fluctuations in physiological pH and other factors can trigger premature drug release in healthy tissues, reducing the amount of drug that reaches the target site. Lastly, challenges in large-scale production may limit the use of pH-dependent polymeric nanoparticles for antiviral drug delivery. Scaling up the production of pH-sensitive nanoparticles while maintaining consistent EE and quality is a significant challenge for industrial applications (87, 88).

To enhance pH-driven encapsulation efficacy, stabilizers can be incorporated judiciously, acidification points optimized precisely, and sophisticated acid-base regulators utilized. These strategies collectively mitigate the impact of constraints on the process. Integrating pH-driven approaches with other encapsulation techniques could potentially lead to improved EE. Combining pH sensitivity with other triggers like redox or enzymatic activity can offer more precise and controlled drug release. Strategies such as incorporating protective shells or surface modifications can enhance nanoparticle stability and prevent premature drug release. Modifying nanoparticle surfaces and utilizing advanced polymers can enhance intracellular drug retention and overcome issues like multidrug resistance (82, 89, 90).

Additionally, when assessing pH-dependent nanoparticle drug delivery, stability is also a potential limitation. Environmental factors like temperature, pH, ionic strength, and solvent type impact micelle stability. Temperature affects micelle formation and structure, with high temperatures potentially disrupting micelles. pH influences charge state and interactions, potentially causing micelles to loosen or change structure. Ionic strength impacts electrostatic interactions, affecting micelle formation and stability. Solvent and additives also play a role in micelle formation. These factors are crucial to consider when interpreting and applying findings, as changing conditions can significantly alter micelle behavior (91).

The instrumentation typically used to assess polymer and nanoparticle characteristics also possess inherent limitations. ^1^H NMR as inherent limitations in characterizing mPEG-PCL copolymers, including difficulty determining average molecular weight and distribution, as well as accurately assessing polydispersity index (PDI). Furthermore, it cannot provide direct information on morphology or particle size of nanoparticles. Consequently, complementary techniques like Gel Permeation Chromatography (GPC) are essential for comprehensive characterization. Scanning electron microscopy (SEM) analysis of mPEG-PCL micelles is limited by sample drying, which can cause micelle collapse or distortion. Additional limitations include low contrast between polymer materials and background, as well as electron beam damage to soft polymer structures. Dynamic Light Scattering (DLS) is useful for characterizing micelle size and polydispersity but has limitations. Detected sizes ranged widely, with a mean size of 63.44 nm and higher-order sizes resulting from aggregation species, corroborated by SEM. However, DLS assumes spherical particles and struggles with polydisperse systems, potentially masking different populations. It is also sensitive to environmental factors like temperature and viscosity, and accuracy is compromised with low concentrations or opaque samples. Therefore, DLS was used in conjunction with other characterization techniques to provide a comprehensive understanding of the micelles and copolymers.

Zeta potential analysis of our mPEG-PCL based nanoparticles showed a near neutral surface charge of −3.28 mV. Near-neutral nanoparticles exhibit extended circulation times due to low surface charge, evading rapid immune clearance and increasing target site accumulation. The mononuclear phagocyte system (MPS) clears foreign particles, but near-neutral particles remain less visible. This is due to the near negligible opsonization that near neutral nanoparticles may undergo. Opsonization occurs when nanoparticles entering the bloodstream become coated with proteins, facilitating immune recognition and clearance. Near-neutral particles minimize opsonin adsorption and clearance due to reduced protein adsorption. This is why PEGylation, which involves coating nanoparticles with polyethylene glycol (PEG), creates a protective shell that blocks protein adsorption.

Our investigation into the efficacy of ribavirin in conjunction with mPEG-PCL pH-dependent, biodegradable micellar nanoparticles (MNPs) demonstrates that this combination significantly attenuates Zika virus replication in JEG-3 placenta carcinoma cells, surpassing the efficacy of free ribavirin administration. The MNPs enhance cellular uptake of ribavirin, augmenting its suppression of Zika virus RNA synthesis and diminishing viral infectivity. Notably, the MNPs mitigate cytotoxicity even at high ribavirin concentrations (12µM) (92). While the findings are promising, limitations include the focus on a single cell line and virus strain, assessment of antiviral efficacy at a single time point, and the use of an in vitro model, which may not accurately reflect in vivo systems. Future studies will investigate the mechanisms underlying the enhanced antiviral activity of ribavirin-loaded MNPs.

Zika virus immunofluorescence (IFA) detection of viral foci is limited to a single cell line (JEG-3 cells) and virus strain (African Zika virus strain MR766), potentially restricting its representativeness. Future studies will investigate the mechanisms underlying the enhanced antiviral activity of ribavirin-loaded MNPs. Our analysis assesses antiviral efficacy at a single time point (3 days post-infection), which may not capture long-term effects. Additionally, the in vitro model used may not accurately reflect in vivo systems, and the reproducibility and variability of MNP preparation require further examination.

The preceding analysis was conducted in an in vitro cell culture environment, which provides a foundational method for evaluating drug-loaded pH-dependent micellar nanoparticles, but possesses associated caveats. Cell culture models lack the intricate cellular interactions and systemic factors inherent to living organisms, and accurately replicating human disease within these models poses considerable challenges (93). Results derived from cell culture studies may not directly translate to the complex environment of the human body. Certain studies have utilized high concentrations of ribavirin, potentially influencing the validity of the findings.

Ribavirin’s immunomodulatory effects are not captured in cell culture settings, potentially influencing its overall efficacy in vivo. Although ribavirin inhibits IMPDH, this inhibition alone may be insufficient to inhibit viral replication in cell culture. Studies investigating ribavirin’s mutagenic effects have yielded conflicting results across different cell culture systems(94). Certain cell lines, such as Vero cells, may exhibit reduced sensitivity to the drug, impacting result interpretation (95).

Overcoming these limitations may require several future studies and considerations. The development of advanced cell culture models could be an asset. Researchers are developing more sophisticated 3D culture models, including organoids and organ-on-chip systems, to better replicate in vivo environments (96). Analyzing viral and cellular changes over time following ribavirin treatment can provide valuable insights into the kinetics of ribavirin’s antiviral effects. Investigating specific mechanisms of action would also be helpful. These studies often employ techniques such as gene expression analysis (e.g., assessing the induction of interferon-stimulated genes or the modulation of key cellular proteins involved in the viral lifecycle) or enzymatic activity assays. Combination studies would also be highly beneficial. Researchers frequently examine the synergistic effects of ribavirin when combined with other antiviral agents, such as interferon-alpha (61). While cell culture studies provide a foundation for understanding ribavirin’s effects, their limitations underscore the importance of utilizing a multifaceted approach to investigate antiviral pharmaceuticals effectively.

The analysis of ribavirin-loaded mPEG-PCL nanoparticles in mice will be a crucial area of research, focusing on efficacy, safety, and pharmacokinetics. The selection of a suitable mouse model susceptible to the targeted disease or condition is essential. The more ideal strains include STAT-1-deficient mice or IFNAR1-deficient mice (e.g. A129 orAG129). Establishing a baseline involves collecting data on body weight, behavior, and relevant physiological parameters prior to treatment. The route of administration and dose determination are critical considerations. Dose-response studies identify the minimum effective dose and maximum tolerated dose, informing the treatment regimen based on ribavirin pharmacokinetics and nanoparticle half-life. Disease monitoring tracks disease progression through clinical scores, viral loads, and biomarkers. Survival studies compare survival rates in treated versus control groups, while tissue analysis, plaque reduction assays, and RT-PCR quantify viral load and assess virus suppression. Blood sampling and tissue distribution analysis elucidate nanoparticle behavior, including distribution, clearance, and potential accumulation in organs. Toxicity evaluation encompasses clinical sign observation, histopathological analysis, and assessment of hematological and biochemical parameters. Immunotoxicity assessment reveals potential effects on the immune system. The final stage involves comparing efficacy and toxicity of ribavirin-loaded nanoparticles with free ribavirin, demonstrating the advantages of the nanoparticle delivery system and modifying properties as necessary to optimize performance. Adherence to ethical guidelines and regulations for animal research is paramount, requiring appropriate approvals prior to conducting experiments.

Nanomaterial development and customization have emerged as a pivotal tool in the design and application of nanomedicine, particularly in the expansion of simple drug delivery systems such as mPEG-PCL micellar nanoparticles. This is exemplified in the assembly of a drug delivery system tailored for the intracellular delivery of ribavirin. The MNPs, comprised of an mPEG-PCL diblock copolymer shell, exhibit the capacity to dissociate at a pH of 5.5, thereby facilitating the intercellular release of the drug cargo. This phenomenon is significant, given that the decrease in endosomal pH is crucial for the internalization of biomolecules and pathogens (97). This dissociation event facilitated the liberation of the antiviral agent, ribavirin, into the cytosolic compartment, as corroborated by ribavirin and Nile Red uptake assays. The findings presented herein indicate that mPEG-PCL copolymer-based micellar nanoparticles not only function as an efficacious ribavirin delivery system but also mitigate the cytotoxic effects associated with free ribavirin, potentially attributable to the enhanced solubility of the pharmaceutical conferred by the nanoparticles, thereby augmenting the internalization of the therapeutic agent by infected cells (98). Further investigation will facilitate the refinement of these methodologies for the targeted inhibition of various human pathogenic viruses, including Ebola, SARS-CoV-2, and mosquito-borne viruses, thereby advancing the field.

Regarding alternative applications, Ribavirin encapsulated in mPEG-PCL drug delivery systems although developed to target Zika virus, have broader applications. These systems offer a platform for delivering ribavirin and potentially other antivirals against Flaviviridae viruses like hepatitis C virus (HCV) and Dengue virus, as well as other viral infections. Beyond viral applications, the system can be used for delivering drugs to treat brain diseases or cancers. The mPEG-PCL drug delivery system, originally developed for Zika virus, has potential for broader use. It can be adapted to target other Flaviviridae viruses like HCV and Dengue virus, as well as be used for delivering drugs to treat brain diseases or cancers. This highlights the versatility and potential of this drug delivery system beyond its initial application. The versatility of this ribavirin-mPEG-PCL system is not relegated to a single virus family, or a single polymer or antiviral.

The development of pH-dependent, biodegradable mPEG-PCL micellar nanoparticles (MNPs) represents a promising approach for the targeted delivery of ribavirin, a broad-spectrum antiviral medication, against Zika virus infections. Our findings demonstrate that ribavirin-loaded MNPs significantly attenuate Zika virus replication in JEG-3 placenta carcinoma cells, surpassing the efficacy of free ribavirin administration. The MNPs enhance cellular uptake of ribavirin, augmenting its suppression of Zika virus RNA synthesis and diminishing viral infectivity while mitigating cytotoxicity. While our results are promising, limitations exist, including the focus on a single cell line and virus strain, assessment of antiviral efficacy at a single time point, and the use of an in vitro model, which may not accurately reflect in vivo systems. Future studies will investigate the mechanisms underlying the enhanced antiviral activity of ribavirin-loaded MNPs and explore the potential applications of this system in delivering antivirals against other viral infections, including Flaviviridae viruses like HCV and Dengue virus. The versatility of PEGylated drug delivery system extends beyond viral applications, with potential uses in treating brain diseases or cancers. Further research is necessary to optimize the design and development of these nanoparticles, addressing challenges such as premature drug release, scalability, and stability. Overall, the ribavirin-mPEG-PCL system holds promise as a cost-effective and readily synthesizable antiviral platform, particularly in resource-limited regions where viral outbreaks pose significant challenges.

## Conclusion

This investigation endeavors to elucidate the antiviral efficacy of ribavirin-loaded mPEG-PCL nanoparticles in mitigating Zika virus infection within cultured JEG-3 cells. Our findings unequivocally demonstrate that this nanoformulation substantially enhances the antiviral activity of ribavirin, resulting in a pronounced reduction in viral load and cytopathic effects in both in vitro and in vivo models compared to the administration of the free drug. The robust antiviral efficacy observed in this study represents a crucial milestone in the development of targeted antiviral therapies for Zika virus, effectively minimizing the systemic toxicity associated with high-dose ribavirin treatment. Notably, our research demonstrates for the first time that mPEG-PCL nanoparticles can be effectively utilized to deliver ribavirin, thereby opening avenues for similar nanomedicine applications against other flaviviruses. The significant potential for clinical translation of this nanoformulation as a more effective and less toxic treatment for Zika virus is underscored by the pronounced antiviral efficacy observed in this study. Furthermore, an important issue emerging from these findings is the potential for repurposing existing antiviral drugs by leveraging advanced nanodelivery systems. The successful inhibition of Zika virus by our formulation provides strong preclinical evidence that warrants further investigation into the use of nanoparticle-encapsulated antivirals for emerging pathogens. Consequently, future research endeavors should focus on elucidating the underlying mechanisms of this nanoformulation and exploring its potential applications in combating various viral infections. The development of nanoparticle-based delivery systems for antiviral medications holds considerable promise for enhancing treatment efficacy while minimizing adverse effects. Regardless of limitations, our findings contribute to the growing body of evidence supporting the potential of nanomedicine in addressing the challenges posed by emerging viral pathogens. The ribavirin-loaded mPEG-PCL nanoparticles investigated in this study demonstrate significant potential as a targeted antiviral therapy for Zika virus, and the successful application of this nanoformulation warrants further investigation into its clinical translation and broader applications in nanomedicine.

## 6. Supplemental Data

**Figures S-1and S-2. Analysis of Nile Red-infused mPEG-PCL MNPs.**

Following the synthesis of the mPEG-PCL copolymer and its subsequent formation into MNPs, Nile Red was incorporated into mPEG-PCL MNPs to analyze the cellular uptake efficiency of these MNPs (see Figures S-1 and S-2). The synthesis of Nile Red-loaded MNPs was conducted following the same protocol used for ribavirin-loaded MNPs. The resulting data is discussed in the Results section of this publication and remained within an acceptable range to facilitate cellular uptake. Our Nile Red-loaded mPEG-PCL MNPs not only met the criteria but also exhibited the capability to enable pH-responsive drug release, resulting in full suppression of Zika Virus infection in cultured cells.

## Acknowledgments

We would like to give many thanks to Andrew Diamanduros (Department of Biology, Georgia Southern University), and Darrin Moore (COSM Core Research Laboratory, Georgia Southern University) for their assistance and critical instrument training. Support for this study was furnished by the NSF (Award Number: 2018774) and Georgia Southern University Research and Service Foundation.

## Supplementary Data Figure Legends

**Figure S-1.**
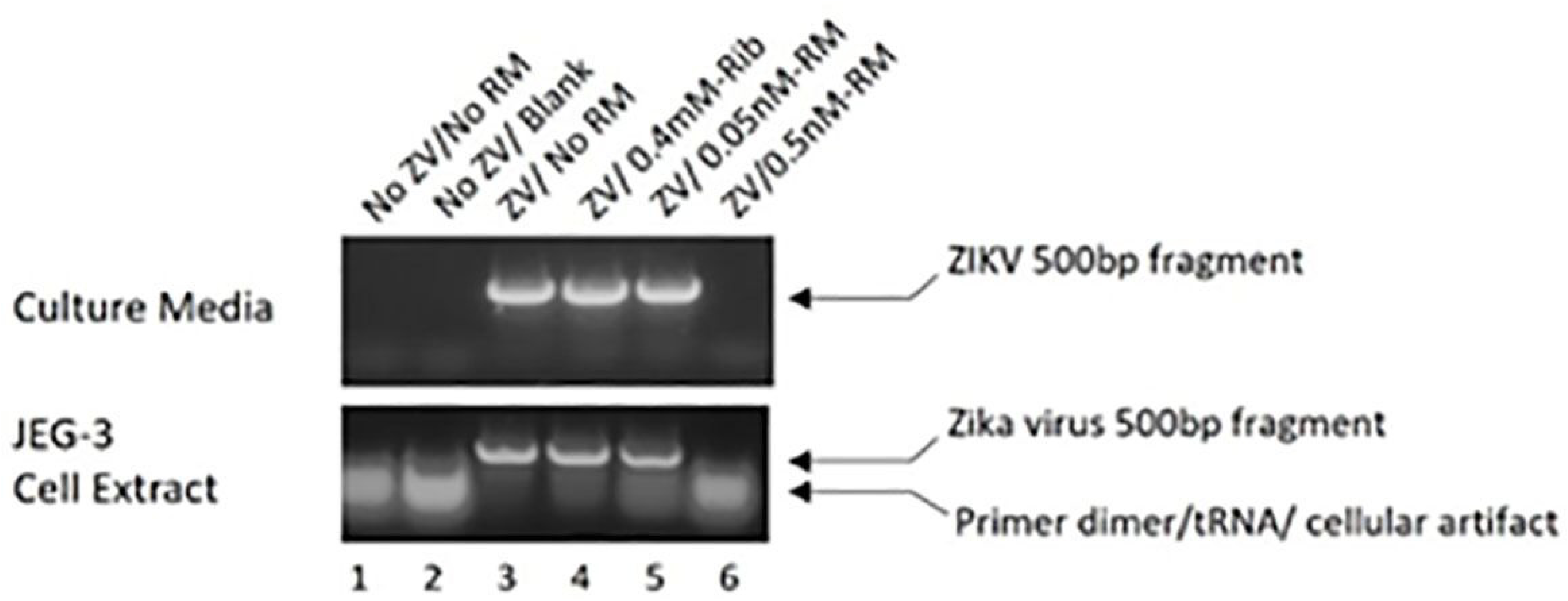
Shown is the mean diameter of our Nile-red loaded mPEG-PCL MNPs by dynamic light scattering (DLS). mPEG-PCL MNPs loaded with Nile-red were monodispersed in a solution possessing KNO_3_ in -pure deionized water.

**Figure S-2.**
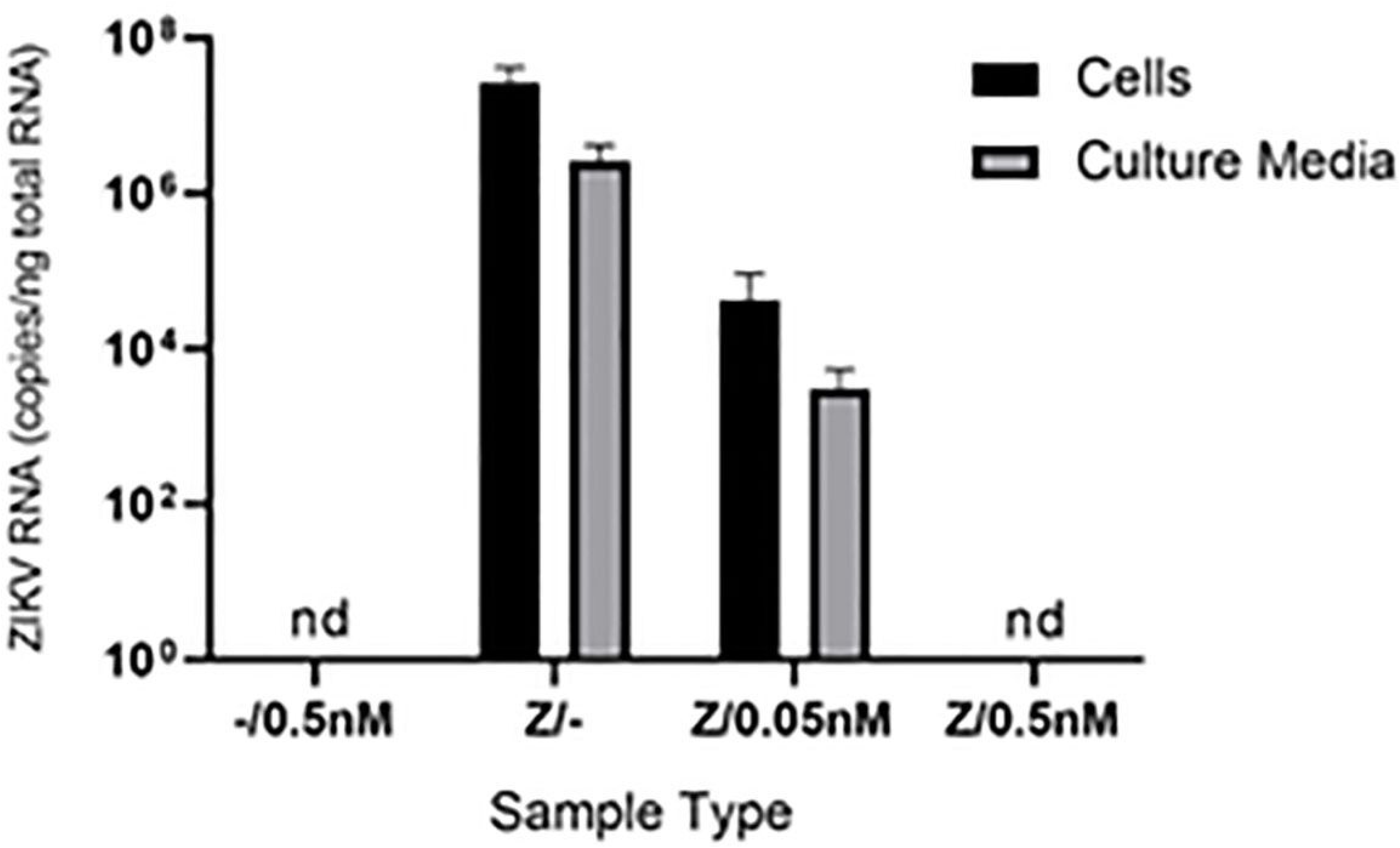
The analysis of both Nile red-loaded and blank mPEG-PCL diblock MNPs was performed by UV-visible spectroscopy was performed on. Each sample was analyzed under ambient conditions.

---

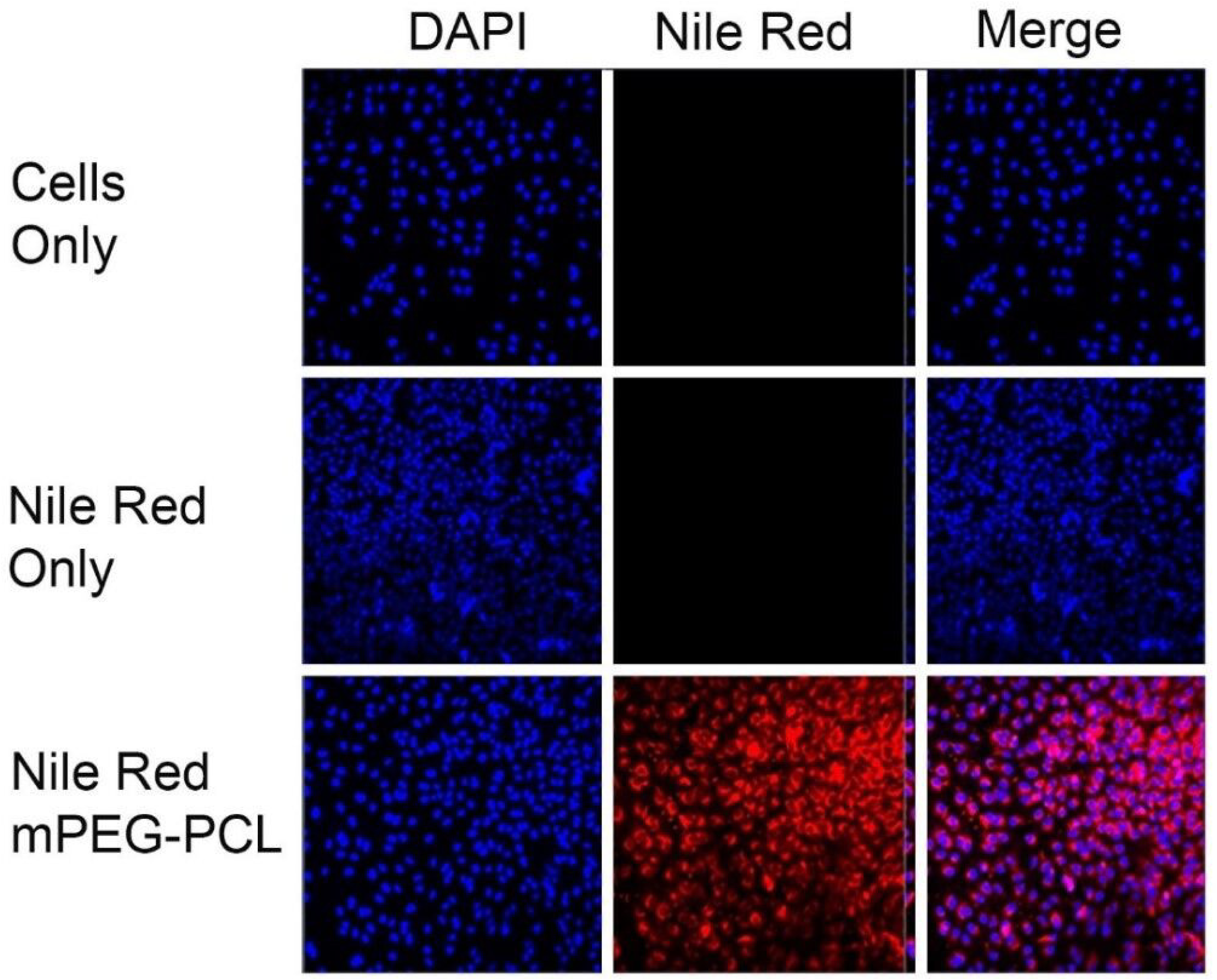

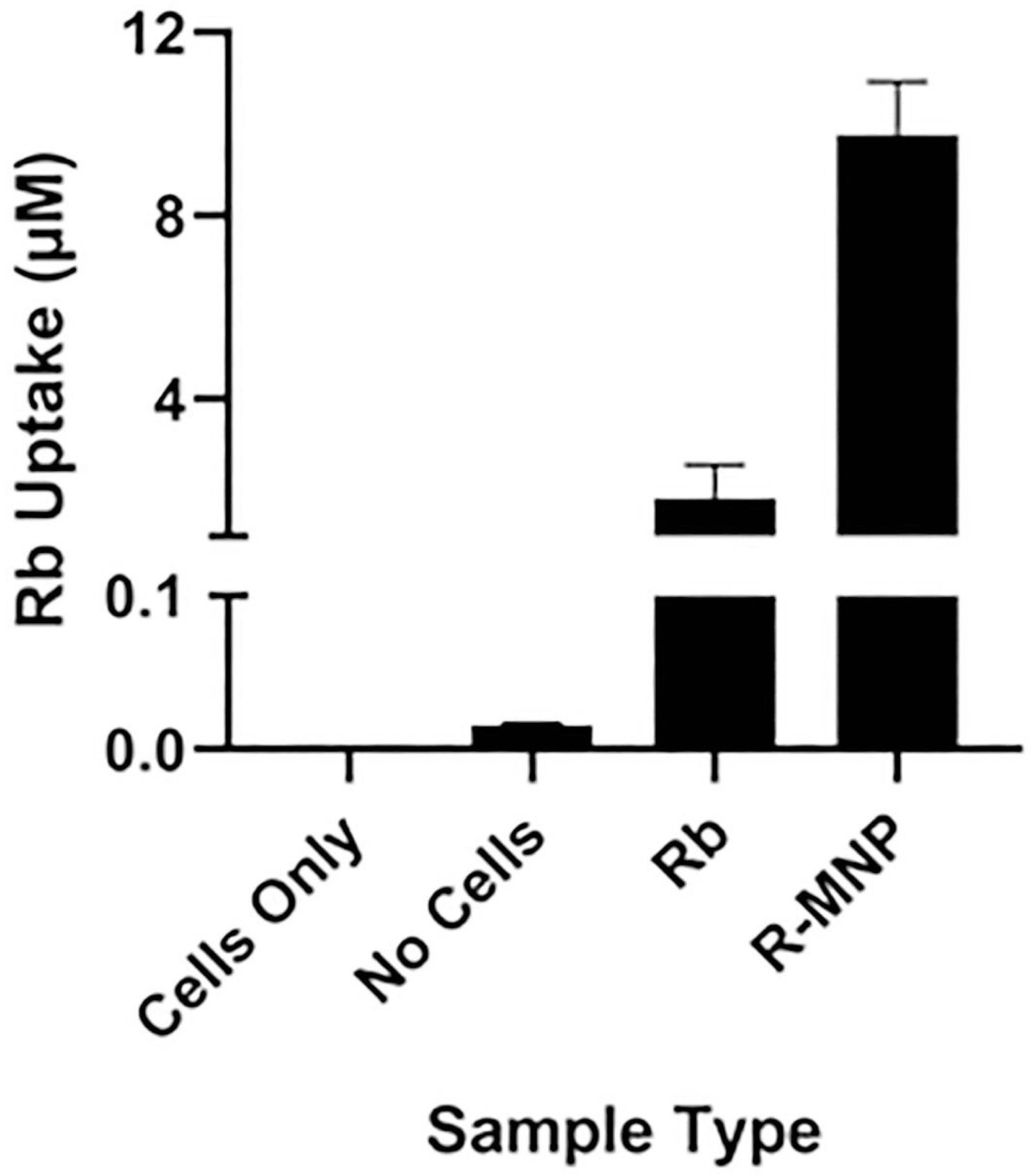

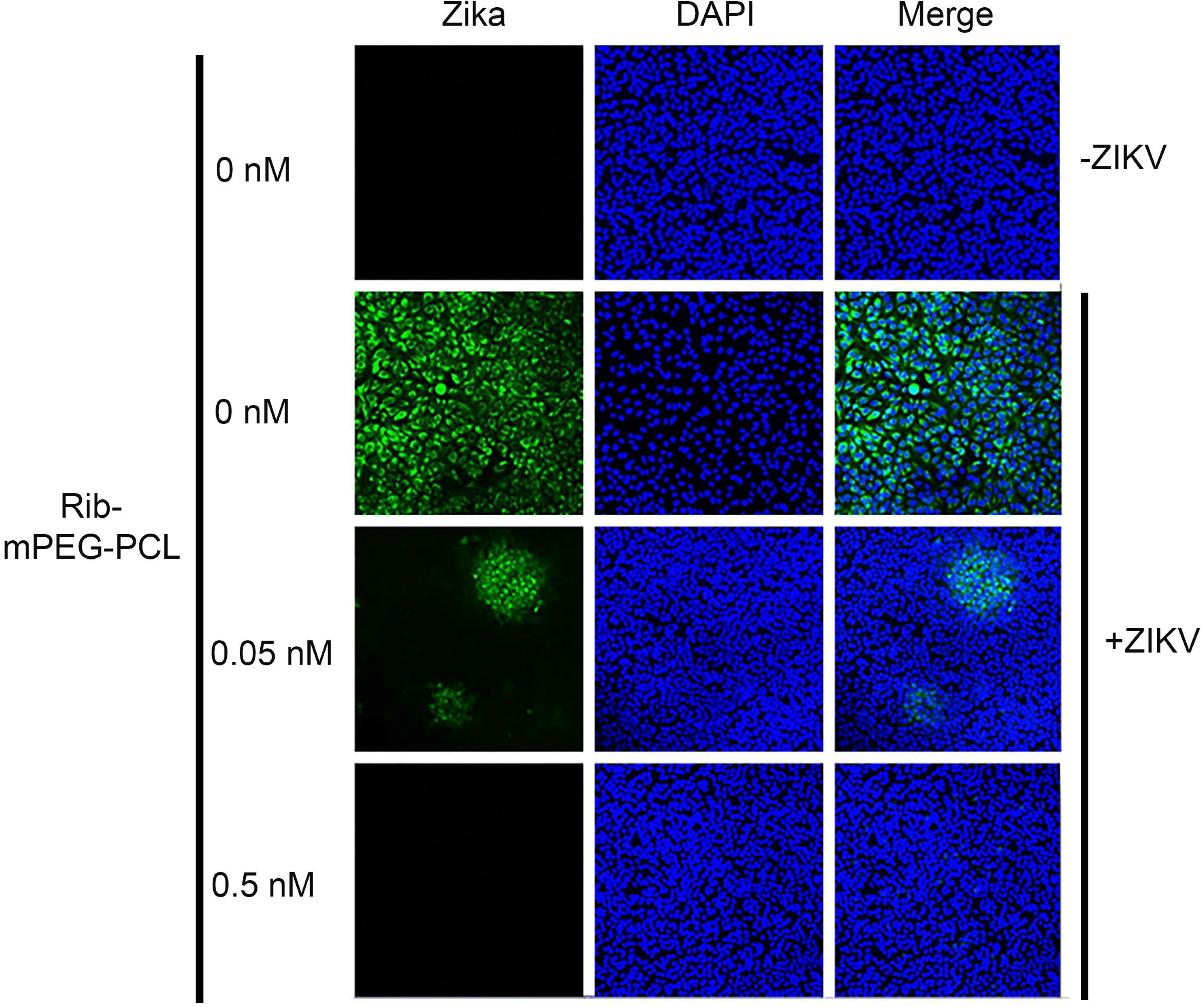

